# PP2A and CDK16 antagonistically regulate WIPI2B phosphorylation and neuronal autophagosome biogenesis

**DOI:** 10.64898/2026.02.12.705597

**Authors:** Heather Tsong, M. Neal Waxham, Andrea K. H. Stavoe

## Abstract

Autophagy is a recycling pathway that clears cellular constituents, supporting homeostasis. In primary murine neurons, autophagosome biogenesis declines during aging. Importantly, this decline can be restored by the ectopic expression of key autophagy component WIPI2B. The phosphorylation state of WIPI2B serine 395 is critical for this restoration, suggesting that WIPI2B S395 phosphorylation regulates autophagosome biogenesis. Here, we identified protein phosphatase 2A (PP2A) and CDK16 as regulators of WIPI2B S395 phosphorylation and neuronal autophagy. Using *Caenorhabditis elegans*, we showed that PP2A and CDK16 regulate neuronal autophagy through the same genetic pathway as WIPI2B *in vivo*. Further, purified mammalian PP2A and CDK16 directly modified WIPI2B S395 phosphorylation *in vitro.* In primary murine neurons, PP2A and CDK16 colocalized with WIPI2B at autophagosomes, and manipulation of PP2A and CDK16 expression altered WIPI2B puncta formation and rates of autophagosome biogenesis. Altogether, our data support the conclusion that PP2A and CDK16 regulate WIPI2B S395 phosphorylation, modulating autophagosome biogenesis in neurons.

## INTRODUCTION

Macroautophagy, hereafter referred to as autophagy, is an evolutionarily conserved cellular recycling pathway used to degrade cellular constituents such as protein aggregates and dysfunctional organelles. Autophagy involves the *de novo* formation of a double membrane vesicle called the autophagosome. The autophagosome subsequently fuses with the endolysosomal system, triggering the degradation of the autophagosomal content (Klionsky and Emr, 2000; Glick et al., 2010).

Autophagosome biogenesis is coordinated by the action of multiple protein complexes. Upon autophagy induction, the initiation complex phosphorylates downstream autophagy components to induce the formation of an autophagosome. The nucleation complex produces phosphatidylinositol 3-phosphate (PI3P) on the initial autophagosomal membrane called the phagophore. PI3P is an important signaling lipid on the phagophore, recruiting key autophagy proteins. Next, the elongation complex contains two conjugation complexes that conjugate ubiquitin-like LC3 to phosphatidylethanolamine on the phagophore, enabling expansion of the phagophore. LC3 is the mammalian ortholog of yeast Atg8 and is essential for phagophore growth. Once the phagophore has engulfed its cargo, the phagophore membrane closes to form a complete autophagosome. The autophagosome then fuses with the endolysosomal system to degrade the contents for reuse (Klionsky and Emr, 2000; Glick et al., 2010; Melia et al., 2020).

Autophagy has historically been studied in yeast and mammalian cell culture, in which acute stressors, notably starvation, trigger massive autophagy induction (Glick et al., 2010; Abada and Elazar, 2014; Hale et al., 2013). However, neuronal autophagy is largely unaffected by these stressors in primary cultures *in vitro*, suggesting that autophagy is differentially regulated in neurons (Tsvetkov et al., 2010; Maday and Holzbaur, 2016). Neurons are highly metabolically active, post-mitotic, and long-lived; as such, neurons heavily rely on autophagy to maintain homeostasis throughout an organism’s lifespan. Neurons have a high basal rate of autophagosome formation compared to non-neuronal cells and autophagosomes are readily delivered to lysosomes for degradation and recycling (Boland et al., 2008). In addition, autophagy is spatially regulated in neurons, with autophagosome biogenesis occurring primarily in the distal axon (Maday and Holzbaur, 2014; Stavoe et al., 2016; Soukup et al., 2016). After formation, cytoplasmic dynein traffics the autophagosome retrogradely along microtubules to the neuronal soma; the autophagosome eventually matures into an autolysosome en route to the soma through fusion with endolysosomes (Maday et al., 2012). The importance of autophagy to neurons is further evidenced by neuron-specific knockout of autophagy components Atg5, Atg7, or Epg5 leading to premature neurodegeneration in mouse models (Hara et al., 2006; Komatsu et al., 2006; Zhao et al., 2013). Additionally, dysregulated autophagy has been implicated in many neurodegenerative diseases (Menzies et al., 2017; Son et al., 2012; Wong and Cuervo, 2010). Together, these data underline the importance of autophagy in neuronal homeostasis and survival.

We previously observed that the rate of autophagosome biogenesis, as monitored by GFP-LC3B puncta formation, decreases significantly with age in primary murine dorsal root ganglia (DRG) neurons (Stavoe et al., 2019). Remarkably, we were able to restore the rate of autophagosome biogenesis in DRG neurons from aged mice to that of young adult mice with the ectopic expression of a single autophagy component, WIPI2B. This restoration of autophagosome biogenesis in neurons from aged mice was dependent on the phosphorylation state of WIPI2B at serine 395 (S395): in DRG neurons from aged mice, phospho-dead WIPI2B(S395A) could restore autophagosome biogenesis while phospho-mimetic WIPI2B(S395E) could not. These data indicate that WIPI2B S395 phosphorylation is a negative regulator of WIPI2B function in autophagosome biogenesis (Stavoe et al., 2019). The kinases and phosphatases that regulate WIPI2B S395 phosphorylation in neurons have not been identified.

WIPI2B is a member of an ancient family of proteins called PROPPINs (beta-propellors that bind to phosphoinositides). There are four PROPPIN family members in mammals: WIPI1 through WIPI4 (**<u>W</u>**D-repeat protein **i**nteracting with **<u>p</u>**hospho**i**nositides) (Proikas-Cezanne et al., 2015). WIPI1 and WIPI2 are related to yeast Atg18, while WIPI3 (also known as WDR45b) and WIPI4 (also known as WDR45) belong to a separate paralogous group related to yeast Ygr223c (Polson et al., 2010). WIPI2B bridges PI3P generation with LC3 conjugation by acting as a scaffold, binding to PI3P on the phagophore and recruiting the conjugation complex through its interaction with conjugation complex member ATG16L1 (Dooley et al., 2014).

Given our previous findings, WIPI2B is a promising target to modulate autophagosome biogenesis, with the goal of upregulating neuronal autophagy to combat aging. Moreover, our data indicate that phosphorylation regulates WIPI2B function, implying the existence of enzymes that regulate that phosphorylation. The regulators of WIPI2B phosphorylation are likely to be better therapeutic targets for modulating WIPI2B function than overexpression of WIPI2B. As such, we sought to identify the molecular regulators of WIPI2B phosphorylation in neurons.

Here, we identified protein phosphatase 2A (PP2A) and CDK16 as regulators of WIPI2B S395 phosphorylation. Using *Caenorhabditis elegans* as an *in vivo* screening platform, we identified PP2A and CDK16 as genetic regulators of neuronal autophagy, likely through their action on WIPI2B S395. We then verified direct enzymatic activity of each protein on mammalian WIPI2B through cell-free *in vitro* phosphorylation assays. We further observed colocalization of both PP2A and CDK16 with WIPI2B and with autophagosomes in primary murine neurons. Additionally, we manipulated expression levels of PP2A and CDK16 in primary murine neurons; we observed that ectopic expression of CDK16 diminished WIPI2B puncta formation while siRNA-mediated knockdown of a PP2A decreased rates of autophagosome biogenesis. Together, our data indicate that PP2A and CDK16 regulate WIPI2B phosphorylation in neurons, thereby modulating autophagosome biogenesis.

## RESULTS

### ATG-18 S375 phosphorylation regulates neuronal autophagy in *C. elegans*

Phylogenetic analysis has shown that the human WIPIs belong to two paralogous groups, with WIPI1 and WIPI2 belonging to one group along with yeast Atg18, and WIPI3 and WIPI4 belonging to another paralogous group with yeast Ygr223c (Polson et al., 2010). In *C. elegans*, each of these groups has only one member; ATG-18 is the nematode ortholog of WIPI1 and WIPI2, and EPG-6 is the ortholog of WIPI3 and WIPI4 (Proikas-Cezanne et al., 2015; Takacs et al., 2019). To use *C. elegans* as a screening platform to identify regulators of WIPI2B S395 phosphorylation, we first asked whether worms have a phosphorylation site in ATG-18 orthologous to WIPI2B S395. ATG-18 has one phosphorylation site, serine 375 (S375), that was experimentally identified through mass spectrometry (Zielinska et al., 2009). Like WIPI2B S395, ATG-18 S375 is located in the disordered C-terminal tail region of the protein (Fig. 1A).

**Figure 1.**
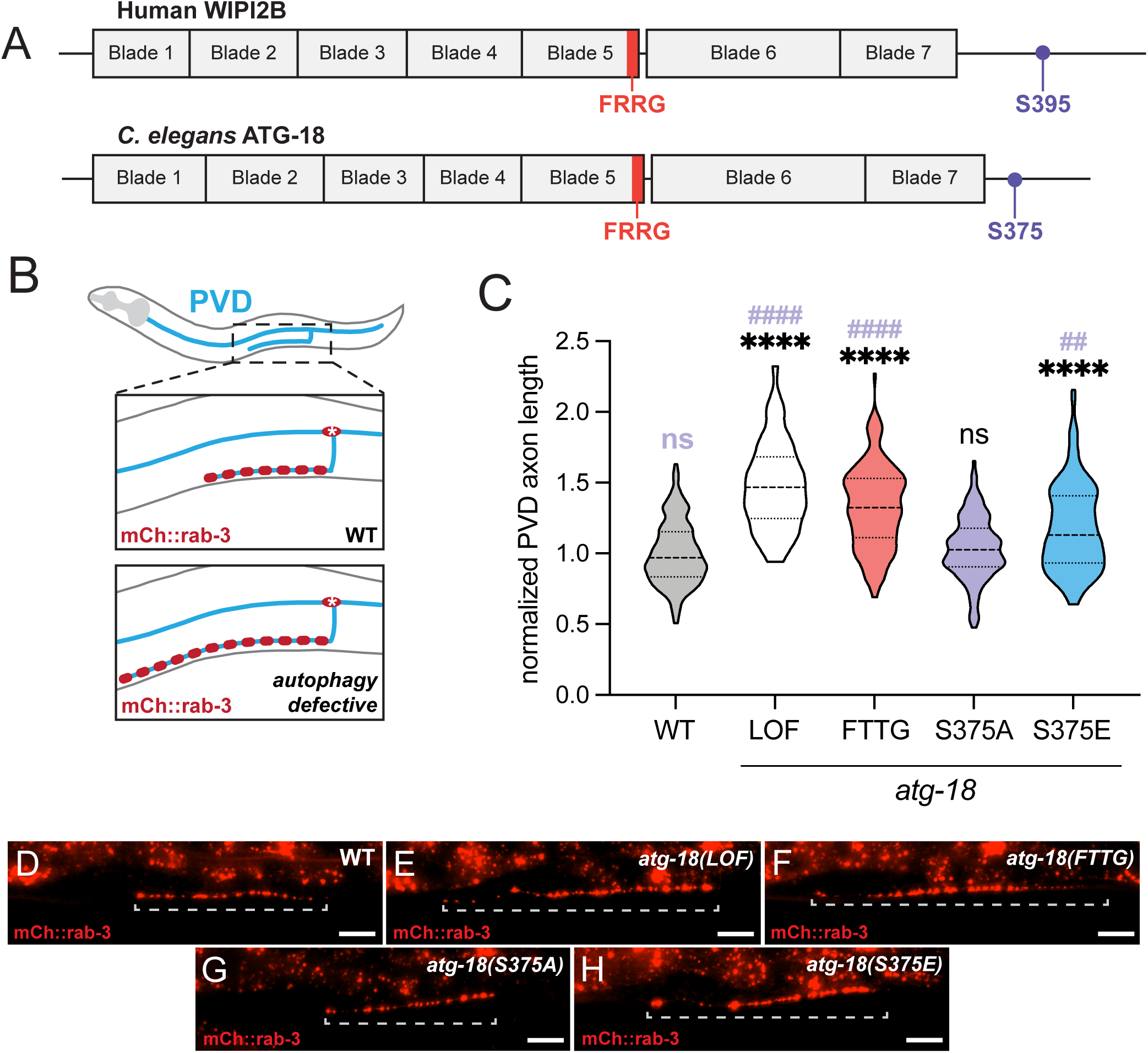
ATG-18 S375 phosphorylation regulates neuronal autophagy in *C. elegans* (A) Schematic of Human WIPI2B and *C. elegans* ATG-18 protein. The structured β-propellor domains are numbered and represented as boxes, while the unstructured regions are represented as a line. The FRRG motif necessary for PI3P binding is indicated in red and the phosphorylation site investigated in this study is in violet. (Β) Diagram of simplified PVD neuron (blue) in *C. elegans*. Schematic of PVD axon (blue) and presynaptic sites (red) in wild-type (top box) and in autophagy mutant animals (bottom box). (C) Quantification of normalized PVD axon length in wild-type and *atg-18* mutant L4 animals (violin plot, median + first and third quartiles; n > 50 animals across three replicates). Comparisons in black are to wild-type and comparisons in purple are to *atg-18(S375A)*; ****/#### p ≤ 0.0001, ## p ≤ 0.01, ns p > 0.05 by Kruskal-Wallis test with Dunn’s multiple comparisons test. (D-H) Representative images of PVD axon presynapses (visualized with mCh::rab-3) in wild-type (D), *atg-18(LOF)* (E), *atg-18(FTTG)* (F), *atg-18(S375A)* (G), and *atg-18(S375E)* (H) L4 animals. Dashed bracket indicates the measured axon. Scale bar: 20 μm.

To interrogate the function of ATG-18 S375 phosphorylation, we used CRISPR to mutate serine 375 at its endogenous locus to either alanine (S375A) or glutamic acid (S375E). Alanine residues cannot be phosphorylated, rendering ATG-18(S375A) constitutively dephosphorylated or phospho-dead. Conversely, the negative charge of glutamic acid is widely used to mimic phosphorylation (Pearlman et al., 2011). Thus, ATG-18(S375E) is considered a phospho-mimetic. As a negative control for ATG-18 function in autophagy, we also generated a lipid-binding deficient mutant. PROPPINs use the conserved FRRG motif to bind to PI3P (Baskaran et al., 2012; Jeffries et al., 2004); previous studies have shown that mutating this motif to FTTG abrogates PROPPIN binding to PI3P and autophagic activity (Dooley et al., 2014; Lu et al., 2011). Thus, we mutated this FRRG motif to FTTG to prevent binding to PI3P, abolishing ATG-18 function in autophagy.

First, to determine if these mutations affected ATG-18 function in neuronal autophagy, we used PVD axon length as a phenotype for impaired neuronal autophagy (Stavoe et al., 2016). PVD is a sensory neuron with a single axon that grows along the ventral nerve cord toward the anterior of the worm. After PVD is born post-embryonically, the PVD axon grows out in a stereotyped manner during larval stages until it reaches the nerve ring by day 1 of adulthood (Smith et al., 2010). At larval stage 4 (L4), the PVD axon has a stereotypic length in wild-type worms (approximately 100 microns). In autophagy mutants, the length of the axon at L4 is significantly longer than that of wild-type worms. This phenomenon is consistent across autophagy mutants representing distinct autophagy complexes, suggesting that autophagy is required to regulate PVD axon outgrowth (Stavoe et al., 2016). Thus, we used the length of the PVD axon at L4 as a readout of neuronal autophagy function, with longer PVD axons correlating with impaired autophagy (Fig. 1B).

We confirmed that ATG-18, as a critical component of the autophagy pathway, phenocopies other loss-of-function autophagy mutants in PVD axon outgrowth. Using a marker for the PVD axon (mCherry::rab-3), we examined PVD axon outgrowth in a putative null allele of *atg-18*. The loss-of-function (LOF) allele *atg-18(gk447069)* contains an ochre nonsense mutation at Q79. In L4 animals, we observed that *atg-18(LOF)* mutants exhibited longer axons than wild-type animals, as expected of an autophagy-defective mutant (Fig. 1C-E). Next, we showed that *atg-18(FTTG)* mutants, in which the PI3P binding motif is mutated, also exhibited longer axons than wild-type animals (Fig. 1C, 1D, 1F), suggesting that ATG-18 does function as expected in autophagy to regulate PVD axon outgrowth.

We then asked how ATG-18 S375 phosphorylation modulated PVD axon outgrowth. The phospho-dead, *atg-18(S375A)*, mutants phenocopied wild-type worms (Fig. 1C, 1D, 1G), suggesting functional autophagy. On the other hand, phospho-mimetic, *atg-18(S375E)*, mutants exhibited longer axons than wild-type worms (Fig. 1C, 1D, 1H), indicative of impaired autophagy. Importantly, the PVD axons of *atg-18(S375E)* animals were also significantly longer than that of *atg-18(S375A)* animals (Fig. 1C, 1G, 1H), highlighting the differential effects of these mutations on PVD axon length. We noted that the effect on PVD axon outgrowth was not as strong in *atg-18(S375E)* mutants as in *atg-18(LOF)* or *atg-18(FTTG)* mutants (Fig. 1C, 1E, 1F, 1H), suggesting that ATG-18 S375 phosphorylation may fine-tune autophagosome formation *in vivo*. Glutamate contains a single negative charge, while actual phosphorylation induces two negative charges (Hunter, 2012). Thus, it is also possible that the glutamic acid is insufficient to fully mimic the effect of ATG-18 S375 phosphorylation on reducing autophagosome biogenesis. Together, our data suggest that the phosphorylation state of ATG-18 S375 regulates neuronal autophagy *in vivo* in *C. elegans,* similar to our previous findings in primary murine neurons.

### PP2A regulatory subunits regulate PVD axon outgrowth in *C. elegans*

We previously showed that ectopic expression of WIPI2B can restore rates of autophagosome formation in neurons from aged mice and that WIPI2B S395 phosphorylation regulates neuronal autophagosome biogenesis (Stavoe et al., 2019). However, the kinases and phosphatases that regulate WIPI2B S395 phosphorylation have yet to be identified. We first focused on identifying a phosphatase that modulates WIPI2B S395. We generated a list of phosphatases that were expressed in neurons and highly conserved across metazoans. We then used LOF mutants of the *C. elegans* orthologs of these phosphatases to determine whether they phenocopied autophagy mutants in PVD axon outgrowth. We hypothesized that if a phosphatase dephosphorylates ATG-18 S375, mutant alleles of that phosphatase would phenocopy *atg-18(S375E)* phospho-mimetic mutants, displaying longer PVD axons at L4 (Fig. 2A).

**Figure 2.**
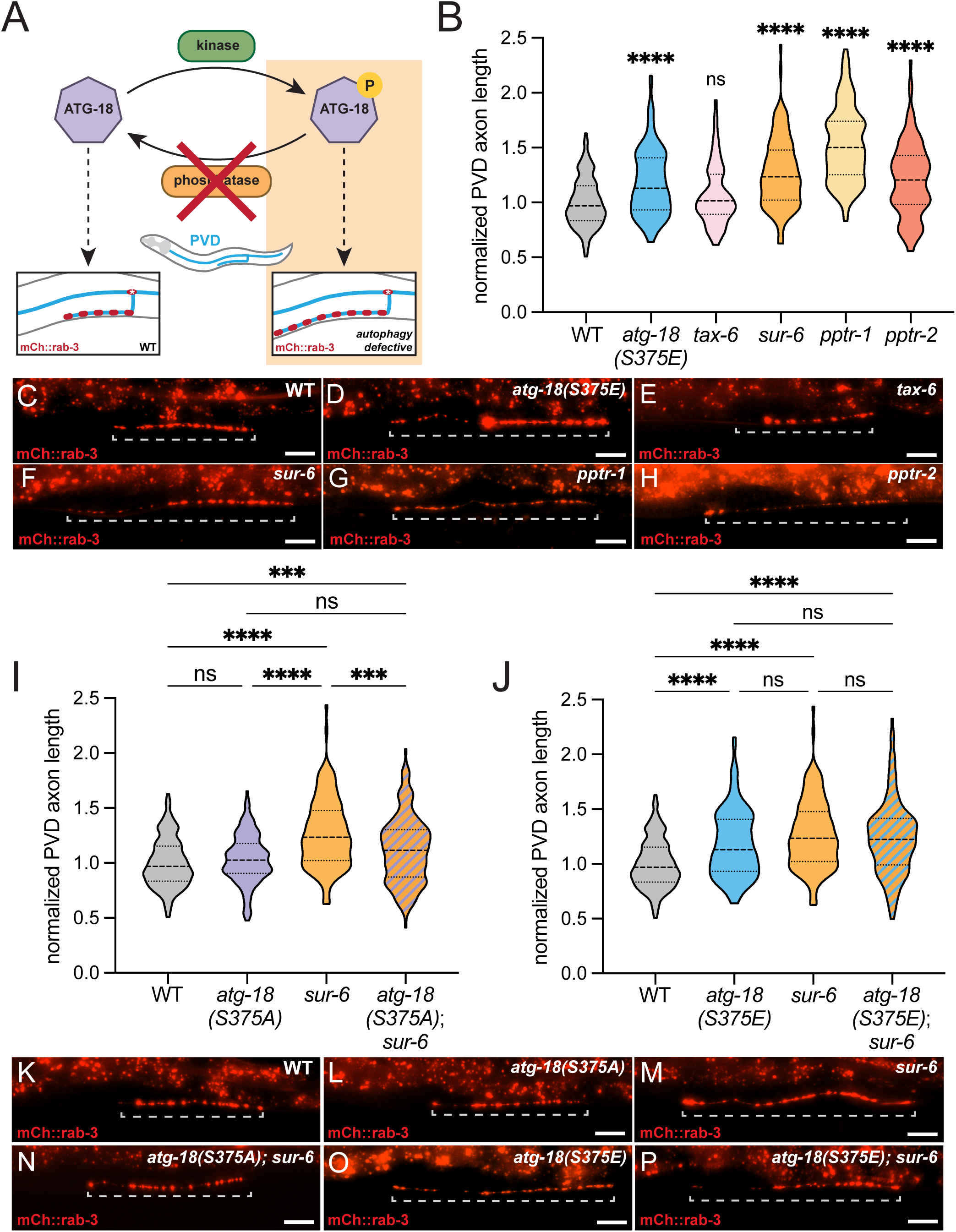
*sur-6* regulates PVD axon length in the same genetic pathway as *atg-18* (A) Diagram summarizing the effect of ATG-18 phosphorylation on the PVD axon phenotype. Light orange box denotes the expected outcome of knocking out the phosphatase. (B) Quantification of normalized PVD axon length in wild-type and phosphatase LOF mutant L4 animals (violin plot, median + first and third quartiles; n > 70 animals across three replicates). Comparisons are to wild-type; ****p ≤ 0.0001, ns p > 0.05 by Kruskal-Wallis test with Dunn’s multiple comparisons test. (C-H) Representative images of PVD axon presynapses (visualized with mCh::rab-3) in wild-type (C), *atg-18(S375E)* (D), *tax-6* (E), *sur-6* (F), *pptr-1* (G), and *pptr-2* (H) L4 animals. Dashed bracket indicates the measured axon. Scale bar: 20 μm. (I-J) Quantification of normalized PVD axon length for epistatic analysis of *sur-6* and *atg-18* in mutant L4 animals (violin plot, median + first and third quartiles; n > 120 across three replicates). ****p ≤ 0.0001, ***p ≤ 0.001, ns p > 0.05 by Kruskal-Wallis test with Dunn’s multiple comparisons test. (K-P) Representative images of PVD axon presynapses (visualized with mCh::rab-3) in wild-type (K), *atg-18(S375A)* (L), *sur-6* (M), *atg-18(S375A); sur-6* (N), *atg-18(S375E)* (O), *atg-18(S375E); sur- 6* (P) L4 animals. Dashed bracket indicates the measured axon. Scale bar: 20 μm.

We first investigated Calcineurin, a serine/threonine protein phosphatase that is modulated by calcium and is highly expressed in neurons (Rusnak and Mertz, 2000). Calcineurin is a Protein Phosphatase 3 (PPP3) family member, and TAX-6 is the worm ortholog of the catalytic subunit of PPP3 (Bye-A-Jee et al., 2020). We found that *tax-6(LOF)* mutant animals had wild-type length PVD axons at L4 (Fig. 2B, 2C, 2E), suggesting that TAX-6/Calcineurin does not regulate neuronal autophagy.

We next examined components of Protein Phosphatase 2A (PP2A). PP2A is a highly conserved and ubiquitous phosphatase that functions as a heterotrimer with catalytic, scaffolding, and regulatory subunits. The regulatory subunit is responsible for subcellular localization and substrate specificity of the complex (Janssens and Goris, 2001; Seshacharyulu et al., 2013; Fowle et al., 2019). PP2A has previously been identified as a regulator of autophagosome biogenesis in cultured mammalian cells (Wong et al., 2015; Fujiwara et al., 2016). In *C. elegans*, mutant alleles of the catalytic subunit, *let-92*, and the PP2A scaffolding subunit, *paa-1*, are lethal (Song et al., 2011). Given that the regulatory subunit provides the target specificity for PP2A activity, we examined alleles of multiple PP2A regulatory subunits. SUR-6 is the worm ortholog of mammalian B55α (PPP2R2A) and B55δ (PPP2R2D). In addition, PPTR-1 and PPTR-2 are worm orthologs of mammalian B56ε (PPP2R5E) and B56δ (PPP2R5D), respectively (Bye-A-Jee et al., 2020). We observed that *sur-6(LOF), pptr-1(LOF),* and *pptr-2(LOF)* mutant animals exhibited longer PVD axons than wild-type animals at L4 (Fig. 2B, 2C, 2F-H), phenocopying *atg-18(S375E)* mutants and suggesting that these genes are involved in neuronal autophagy regulation.

The quantitatively similar phenotypes of the PP2A regulatory subunit mutants and *atg-18(S375E)* mutants are consistent with our hypothesis that PP2A regulates ATG-18 S375 phosphorylation to modulate neuronal autophagosome formation. We next employed epistasis experiments to determine if *sur-6* and *atg-18* function in the same genetic pathway to regulate PVD axon length. If *atg-18* acts downstream from *sur-6* in the same pathway, *atg-18(S375A); sur-6(LOF)* double mutants would have shorter axons than *sur-6(LOF)* single mutant animals. Indeed, we observed that the *atg-18(S375A); sur-6(LOF)* double mutants had shorter axons than the *sur-6(LOF)* single mutant (Fig. 2I, 2M, 2N), indicating that the *atg-18(S375A)* mutation masked the effect of the *sur-6(LOF)* mutation and that *atg-18* is epistatic to *sur-6.* However, *atg-18(S375A); sur-6(LOF)* double mutants had significantly longer PVD axons than wild-type worms (Fig. 2I, 2K, 2N), suggesting that SUR-6 may regulate the phosphorylation state of other residues that themselves modulate autophagosome biogenesis. Additionally, we examined *atg-18(S375E); sur-6(LOF)* double mutants, asking whether losing SUR-6 function in the background of phospho-mimetic ATG-18 would have additive effects. We found that *atg-18(S375E); sur-6(LOF)* double mutants had PVD axons that were not longer than either *sur-6(LOF)* or *atg-18(S375E)* single mutants (Fig. 2J, 2M, 2O, 2P), suggesting that *sur-6* and *atg-18* do not operate in synergistic pathways to regulate PVD axon outgrowth. Together, our data indicate that *atg-*18 acts downstream from *sur-6* in the same genetic pathway and support the conclusion that PP2A dephosphorylates ATG-18 S375 to regulate neuronal autophagy.

### PP2A dephosphorylates WIPI2B *in vitro*

Based on our genetic evidence, we next asked whether mammalian PP2A can directly dephosphorylate mammalian WIPI2B. We performed *in vitro* dephosphorylation assays with purified recombinant human PP2A. As a substrate, we expressed and purified a peptide containing the last 78 amino acids of human WIPI2B, flanked by GST- and HA- tags, termed the WIPI2B tail. Importantly, the WIPI2B tail includes the S395 phosphorylation site (Fig. 3A). We co-expressed Lambda phosphatase in *E. coli* with the WIPI2B tail to limit phosphorylation prior to purification. We quantified changes to WIPI2B phosphorylation via western blot using a phospho-specific antibody for WIPI2B S395. Using a total WIPI2B antibody, we detected a 38 kDa band on the immunoblot, corresponding to the predicted size of the purified WIPI2B tail (Fig. 3B). This band was also faintly labelled by the phospho-specific WIPI2B antibody, yielding a low phospho-WIPI2B/total-WIPI2B ratio (Fig. 3B, 3C).

**Figure 3.**
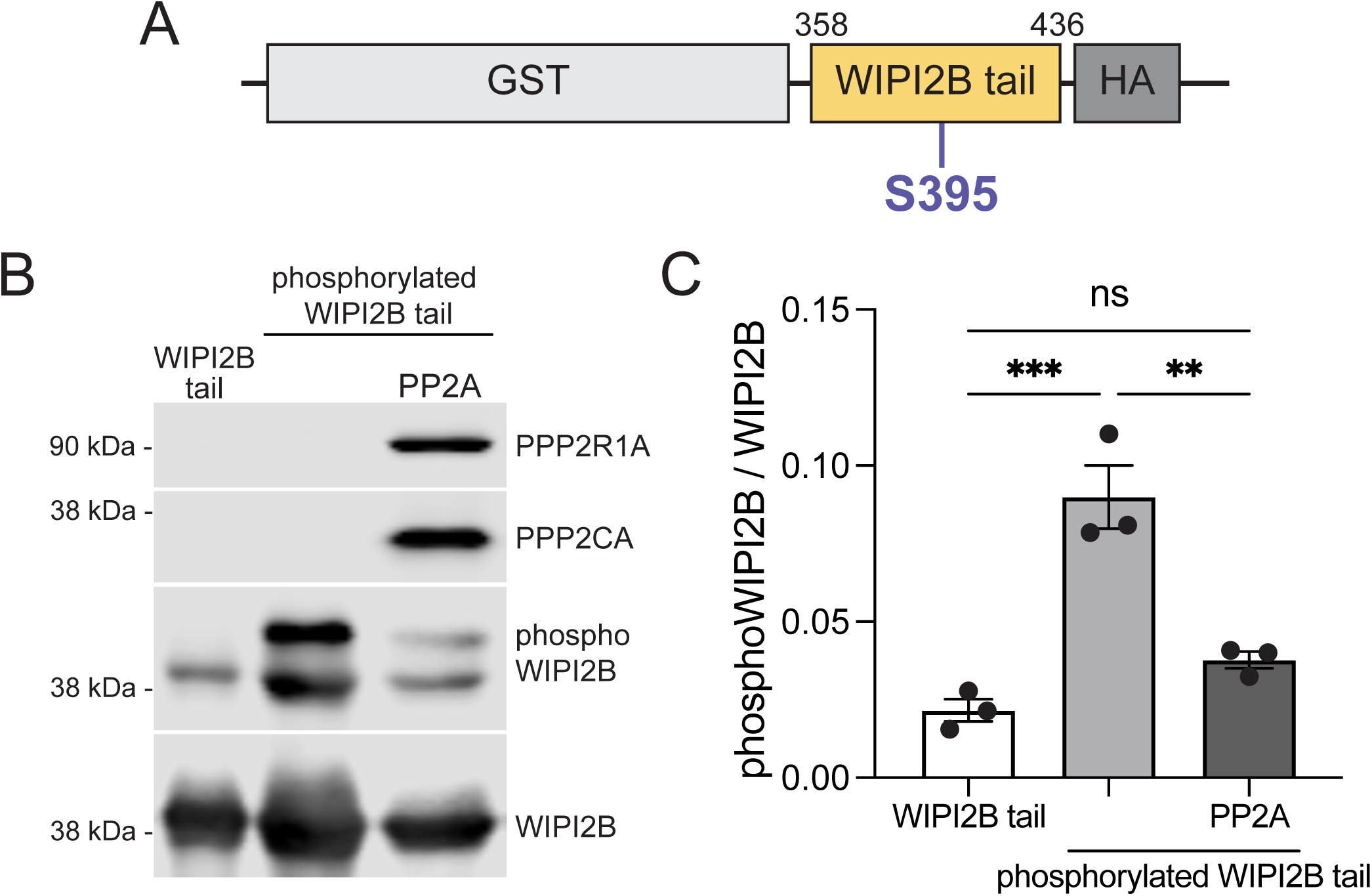
Mammalian PP2A dephosphorylates WIPI2B tail *in vitro* (A) Schematic of the GST-and HA-tagged WIPI2B tail peptide containing the last 78 amino acids of human WIPI2B (amino acids 358-436). The S395 phosphorylation site is indicated in violet. (B) Representative immunoblots of the *in vitro* dephosphorylation assay with purified WIPI2B tail and phosphorylated WIPI2B tail, with and without recombinant human PP2A. (C) Quantification of phosphoWIPI2B signal relative to total WIPI2B signal for the *in vitro* dephosphorylation assay represented in B (mean ± SEM, n = 3). ***p ≤ 0.001, **p ≤ 0.01, ns p > 0.05 by one-way ANOVA with Tukey’s multiple comparisons test.

We hypothesized that whole mouse brain lysate contained kinases that could phosphorylate WIPI2B S395, as we previously detected phospho-WIPI2B in mouse whole brain lysate (Stavoe et al., 2019). Since the purified WIPI2B tail displayed low levels of phosphorylation, we induced phosphorylation by incubating the purified WIPI2B tail with a cytosolic extract of whole mouse brain homogenate. This treatment significantly increased the phosphorylation of the WIPI2B tail as measured by the increased phospho-WIPI2B/WIPI2B ratio (Fig. 3B, 3C). Of note, the phosphorylation treatment led to the appearance of an additional band of higher molecular weight that was strongly labelled by the phospho-specific WIPI2B antibody (Fig. 3B).

Finally, we asked whether PP2A could directly dephosphorylate WIPI2B S395. After washing out the brain homogenate, we then incubated the phosphorylated WIPI2B tail with purified recombinant PP2A. The recombinant human PP2A contained the catalytic PPP2CA subunit and the scaffolding PPP2R1A subunit. Addition of recombinant PP2A to the phosphorylated WIPI2B tail led to a significant reduction in WIPI2B S395 phosphorylation (Fig. 3B, 3C), demonstrating that mammalian PP2A can directly dephosphorylate mammalian WIPI2B S395. Further, our mammalian *in vitro* data are consistent with and confirm our findings *in vivo* in *C. elegans*.

### CDK16 regulates PVD axon outgrowth in *C. elegans*

Next, we sought to identify a kinase that modulates WIPI2B S395 phosphorylation. We again took a candidate approach, generating a list of candidates by cross-referencing a kinase prediction algorithm (PhosphoNET) and a recently published data set of the human serine/threonine phosphoproteome (Johnson et al., 2023). With these tools, we identified fifty candidate kinases that may modulate WIPI2B S395 phosphorylation (Supplementary table S1). We further refined this list by focusing on highly conserved kinases that are expressed in neurons. Once again, we utilized the *C. elegans* PVD axon length phenotype to screen for kinases that regulate neuronal autophagy.

Our evidence indicated that PP2A can modulate WIPI2B S395 phosphorylation. Thus, we reasoned that a kinase that also modulates WIPI2B S395 phosphorylation would oppose PP2A in PVD axon outgrowth. Therefore, the mutant allele of such a kinase would suppress the *sur-6(LOF)* phenotype of long PVD axons, and kinase(LOF); *sur-6(LOF)* double mutants would display wild-type PVD axons. We screened LOF mutant alleles of *pct-1, jnk-1, pmk-1, cdk-5, cdk-1, and unc-43* with *sur-6(LOF)* in the genetic background for the PVD axon length phenotype (see supplementary table S2 for allele information). We found that *pct-1(LOF); sur-6(LOF)* double mutants*, cdk-5(LOF); sur-6(LOF)* double mutants, and *unc-43(LOF); sur-6(LOF)* double mutant animals suppressed the *sur-6(LOF)* phenotype and had PVD axons that were similar in length as wild-type PVD axons (Fig. 4A, 4B, 4D, 4G, 4I), suggesting that these kinases may function in opposition to SUR-6/PP2A to regulate neuronal autophagy. In contrast, *jnk-1(LOF); sur-6(LOF)* double mutants*, pmk-1(LOF); sur-6(LOF)* double mutants, and *cdk-1(LOF); sur-6(LOF)* double mutant animals did not suppress the *sur-6(LOF)* phenotype and had PVD axons that were significantly longer than wild-type PVD axons (Fig. 4A, 4B, 4E, 4F, 4H). These data suggest that these kinases do not oppose SUR-6/PP2A function in neuronal autophagy; therefore, we eliminated these candidates from subsequent experiments.

**Figure 4.**
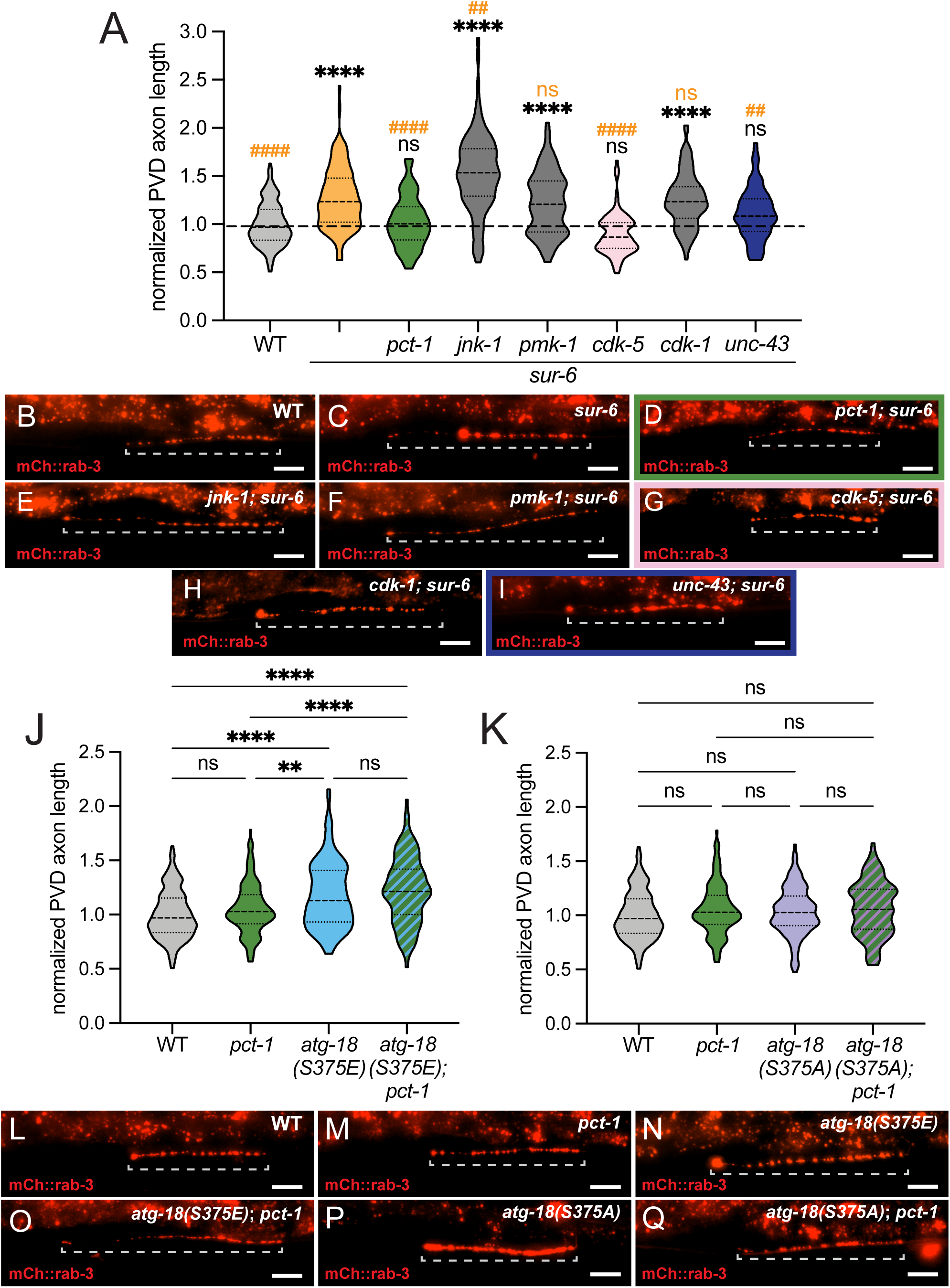
*pct-1* regulates PVD axon length in the same genetic pathway as *atg-18* (A) Quantification of normalized PVD axon length in wild-type and kinase(LOF); *sur-6* double mutant L4 animals (violin plot, median + first and third quartiles; n > 65 animals across three replicates). The dashed line denotes the median for wild-type animals. Comparisons in black are to wild-type and comparisons in orange are to *sur-6(LOF)*; ****/#### p ≤ 0.0001, ## p ≤ 0.01, ns p > 0.05 by Kruskal-Wallis test with Dunn’s multiple comparisons test. (B-I) Representative images of PVD axon presynapses (visualized with mCh::rab-3) in wild-type (B), *sur-6* (C), *pct-1; sur-6* (D), *jnk-1; sur-6* (E), *pmk-1; sur-6* (F), *cdk-5; sur-6* (G), *cdk-1; sur-6* (H), and *unc-43; sur-6* (I) L4 animals. Dashed bracket indicates the measured axon. Scale bar: 20 μm. (J-K) Quantification of normalized PVD axon length for epistatic analysis of *pct-1* and *atg-18* in mutant L4 animals (violin plots, median + first and third quartiles, n > 95 across three replicates). ****p ≤ 0.0001, **p ≤ 0.01, ns p > 0.05 by Kruskal-Wallis test with Dunn’s multiple comparisons test. (L-Q) Representative images of PVD axon presynapses (visualized with mCh::rab-3) in wild-type (L), *pct-1* (M), *atg-18(S375E)* (N), *atg-18(S375E); pct-1* (O), *atg-18(S375A)* (P), and *atg-18(S375A); pct-1* (Q) L4 animals. Dashed bracket indicates the measured axon. Scale bar: 20 μm.

As before, we next employed epistasis experiments to determine if *pct-1*, *cdk-5*, or *unc-43* function in the same genetic pathway as *atg-18* to regulate PVD axon length. If ATG-18 acts downstream from a kinase in the same pathway, we expected double mutants with *atg-18(S375E)* to phenocopy *atg-18(S375E)* single mutant animals. We observed that *pct-1(LOF); atg-18(S375E)* double mutants phenocopied *atg-18(S375E)* single mutant animals (Fig. 4J, 4N, 4O). On the other hand, *cdk-5(LOF); atg-18(S375E)* double mutants and *unc-43(LOF); atg-18(S375E)* double mutants had significantly shorter PVD axons than *atg-18(S375E)* single mutant animals (Fig. S1). These data indicate that the *atg-18(S375E)* mutation masked the effect of the *pct-1(LOF)* mutation, but not of the *cdk-5(LOF) or unc-43(LOF)* mutations and suggest that *atg-18* is epistatic to *pct-1*, but not *cdk-5* or *unc-43*. Additionally, we examined *atg-18(S375A); pct-1(LOF)* double mutant animals and observed wild-type PVD axon length (Fig. 4K, 4L, 4Q), suggesting that *atg-18* and *pct-1* do not function in synergistic pathways to regulate PVD axon outgrowth. Altogether, our data indicate that *pct-1* and *atg-18* function in the same genetic pathway and support the conclusion that PCT-1 phosphorylates ATG-18 S375 to regulate neuronal autophagy *in vivo*.

### CDK16 phosphorylates WIPI2B *in vitro*

PCT-1 is the worm ortholog of mammalian CDK16 (PCTAIRE1). CDK16 is a highly conserved serine/threonine kinase that belongs to a subfamily of cyclin-dependent kinases along with CDK17 and CDK18 (PCTAIRE2 and PCTAIRE3, respectively). These kinases are highly expressed in post-mitotic tissues, including the brain (Cole, 2009; Besset et al., 1999; Hirose et al., 1997; Herskovits and Davies, 2006; de Oliveira Pepino et al., 2021). Due to their expression pattern, many have postulated a role for these kinases outside of the cell cycle; however, the functions and substrates of these kinases are relatively unknown. The most well-studied of this group, CDK16, has roles in a variety of cell processes, including autophagy and neurite outgrowth (Dohmen et al., 2020; Graeser et al., 2002; Ou et al., 2010).

Once again, we utilized the *in vitro* phosphorylation assay with the WIPI2B tail. We already knew that we could induce phosphorylation of the WIPI2B tail through incubation with mouse brain homogenate (Fig. 3B, 3C), implying that a kinase that phosphorylates WIPI2B S395 is present in the whole brain lysate. We reasoned that inhibiting this kinase would prevent WIPI2B phosphorylation. We chose a panel of inhibitors that act on three independent kinases. Cdk1/2 inhibitor III is a highly potent inhibitor of CDK1 and CDK2 activity and has been experimentally verified to inhibit CDK16 kinase activity (Dixon-Clarke et al., 2017). Torin 1 is a widely used inhibitor for mTOR activity (Thoreen et al., 2009). Autocamtide-2-related inhibitor peptide (AIP) is a specific inhibitor for CaMKII activity (Ishida et al., 1995). We incubated the purified WIPI2B tail with mouse brain homogenate, ATP, and each of the kinase inhibitors in separate reactions. We then examined the levels of WIPI2B S395 phosphorylation as quantified by the phospho-WIPI2B/WIPI2B ratio via western blot. We found that addition of AIP to the mouse brain homogenate and purified WIPI2B tail did not significantly diminish the phosphorylation of the WIPI2B S395 (Fig. 5A, 5B). These data suggest that CaMKII does not phosphorylate WIPI2B and support our *in vivo C. elegans* results that eliminated the CaMKII ortholog, UNC-43, as a candidate kinase. Addition of Torin 1 modestly diminished WIPI2B S395 phosphorylation (Fig. 5A, 5B). Finally, addition of Cdk1/2 inhibitor III most strongly inhibited the phosphorylation of the WIPI2B S395 by the mouse brain homogenate (Fig. 5A, 5B). These data suggest that a kinase targeted by the Cdk1/2 inhibitor III can modulate WIPI2B S395 phosphorylation.

**Figure 5.**
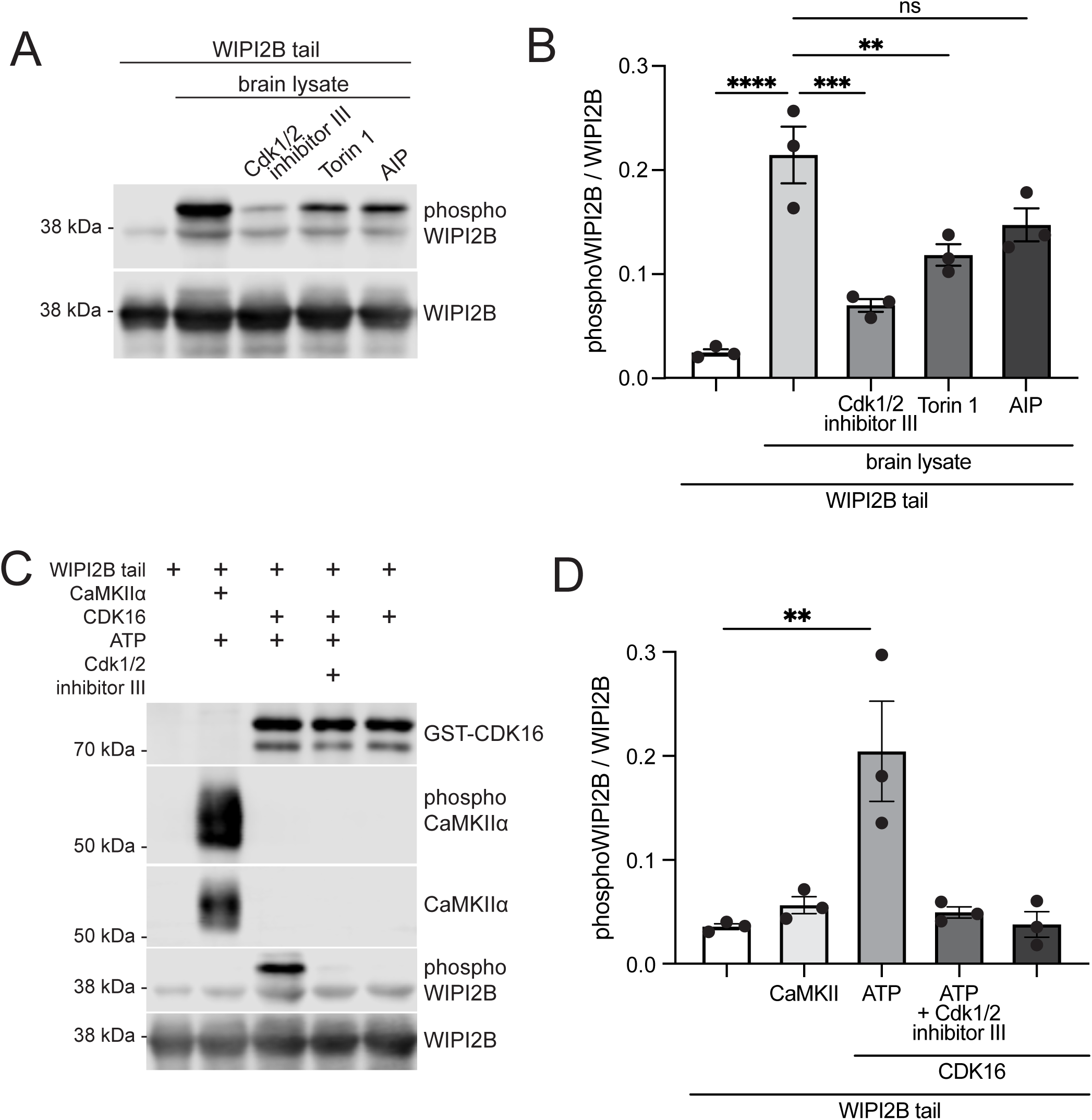
Mammalian CDK16 phosphorylates WIPI2B tail *in vitro* (A) Representative immunoblots of the *in vitro* phosphorylation assay with purified WIPI2B tail treated with mouse brain homogenate and kinase inhibitors. (B) Quantification of phosphoWIPI2B signal relative to total WIPI2B signal for the *in vitro* phosphorylation assay represented in A (mean ± SEM, n = 3). ****p ≤ 0.0001, ***p ≤ 0.001, **p ≤ 0.01, ns p > 0.05 by one-way ANOVA with Tukey’s multiple comparisons test. (C) Representative immunoblots of the *in vitro* phosphorylation assay with purified WIPI2B tail, recombinant CaMKII, and recombinant CDK16. (D) Quantification of phosphoWIPI2B signal relative to total WIPI2B signal for the *in vitro* phosphorylation assay represented in C (mean ± SEM, n = 3). **p ≤ 0.01 by one-way ANOVA with Tukey’s multiple comparisons test.

As CDK16 is a verified target of this inhibitor, we next performed the *in vitro* phosphorylation assay with human recombinant CDK16 and the purified WIPI2B tail. In the presence of ATP and recombinant CDK16 and recombinant Cyclin Y, we detected a significant increase in the phosphorylation of WIPI2B S395 (Fig. 5C, 5D). We did not detect an increase in WIPI2B S395 phosphorylation in reactions containing Cdk1/2 inhibitor III or in reactions without ATP added (Fig. 5C, 5D). As a negative control, we tested whether mammalian recombinant CaMKIIα could phosphorylate the WIPI2B S395 *in vitro*. Auto-phosphorylation of CaMKIIα at T286 indicated that the recombinant CaMKIIα was active and capable of phosphorylation (Fig. 5C). The results of both the *C. elegans* screen and the previous phosphorylation assay with kinase inhibitors indicated that CaMKII does not modulate WIPI2B S395 phosphorylation. In line with these results, we did not detect an increase in WIPI2B S395 phosphorylation when the purified WIPI2B tail was incubated with recombinant CaMKIIα (Fig. 5C, 5D), suggesting that CaMKIIα could not phosphorylate WIPI2B S395 *in vitro*. These data indicate that CDK16, specifically, can directly phosphorylate WIPI2B S395 *in vitro*, supporting the conclusion that CDK16 modulates WIPI2B S395 phosphorylation.

### PP2A and CDK16 regulate autophagosome biogenesis and WIPI2B puncta formation in cultured mammalian neurons

Together, our data indicate that PP2A and CDK16 modulate WIPI2B S395 phosphorylation and suggest that they regulate neuronal autophagosome biogenesis. To directly examine their relative impact on autophagosome biogenesis, we examined their localization and functional effects in primary murine dorsal root ganglion (DRG) neurons expressing the GFP-LC3B reporter for autophagosomes. We first asked whether PP2A, CDK16, and WIPI2B colocalize at autophagosomes. We ectopically expressed Halo-B55α, Halo-B55δ, Halo-CDK16, Halo-CDK17, or Halo-CDK18 along with SNAP-WIPI2B in GFP-LC3B primary DRG neurons and used multi-channel time-lapse spinning disk confocal microscopy to determine colocalization of each of the Halo-labelled constructs with SNAP-WIPI2B and GFP-LC3B at axonal distal tips. In our DRG neuron cultures, we observed cytosolic GFP-LC3B throughout the neurite with ring-like and punctate structures concentrated at axonal tips, consistent with previous descriptions of GFP-LC3B localization in DRG neurons (Maday et al., 2012; Stavoe et al., 2019). Each of the ectopically expressed constructs displayed punctate structures visible above the background cytoplasmic signal (Fig. 6A-E, Fig. S2). We observed colocalization of Halo-B55α and Halo-B55δ puncta with SNAP-WIPI2B puncta (Fig. S2A, S2B). We also observed instances of colocalization between Halo-B55α/δ puncta, SNAP-WIPI2B puncta, and GFP-LC3B puncta (Fig. 6A, 6B). These colocalization events were transient and usually did not persist for the whole 10-min imaging window, consistent with our previous quantifications of WIPI2B transiently residing at developing autophagosomes (Stavoe et al., 2019). Similarly, we observed colocalization of Halo-CDK16, Halo-CDK17, and Halo-CDK18 with SNAP-WIPI2B puncta (Fig. S2C-E) as well as Halo-CDK16/17/18 and SNAP-WIPI2B together with GFP-LC3B (Fig. 6C-E). These data indicate that B55α, B55δ, CDK16, CDK17, and CDK18 can localize with WIPI2B at LC3B-labelled autophagic vesicles, implying that they may function together.

**Figure 6.**
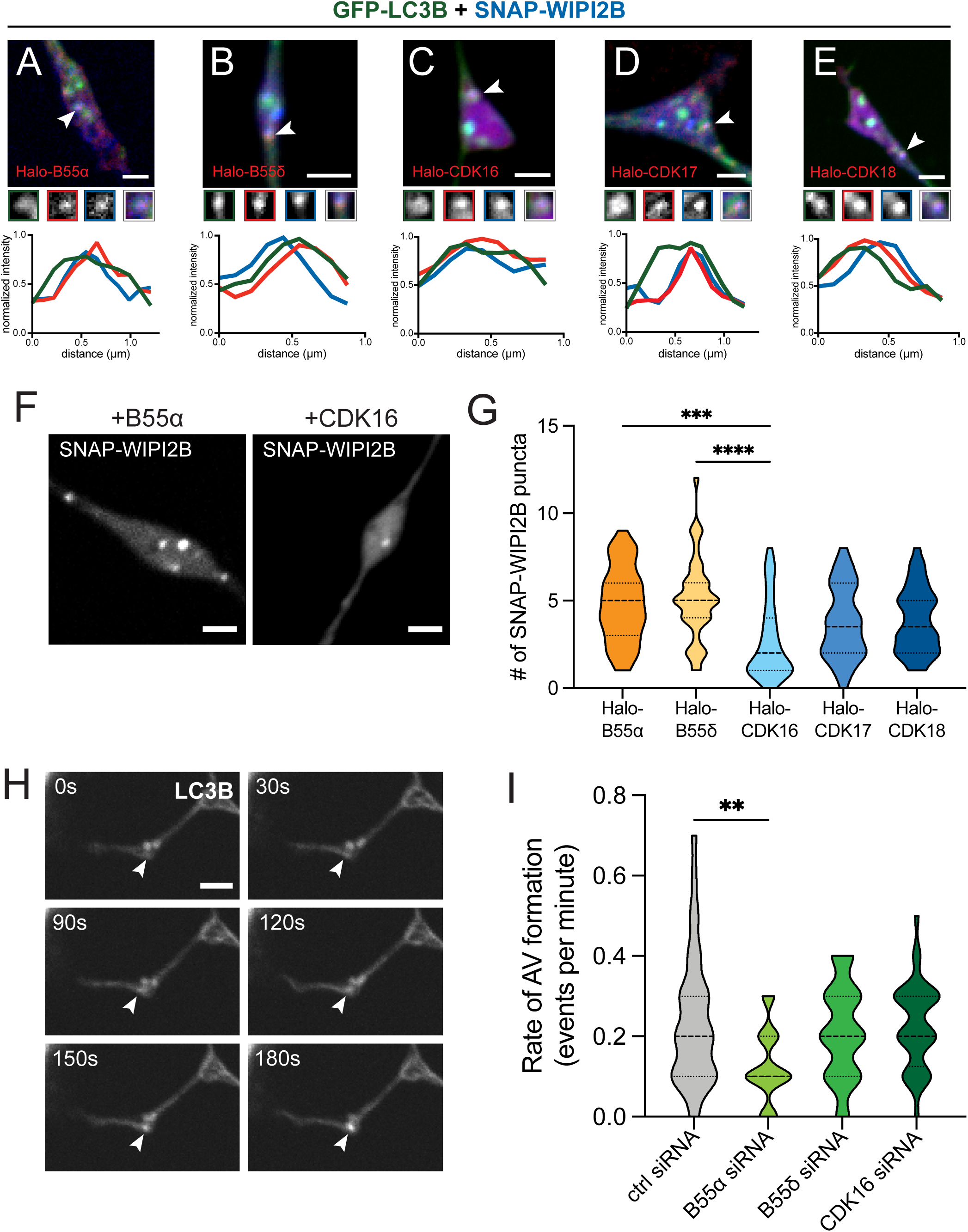
PP2A and CDK16 regulate autophagosome biogenesis and WIPI2B puncta formation. (A-E) Representative merged micrographs of distal axonal tips of GFP-LC3B DRG neurons ectopically expressing SNAP-WIPI2B and Halo-B55α (A), Halo-Β55δ (B), Halo-CDK16 (C), Halo-CDK17 (D), or Halo-CDK18 (E). White arrowheads indicate the colocalization event depicted in the magnified views and quantified in the corresponding line scan plot below the full micrograph. Borders of the magnified views and color of lines in the line scan plot denote the channel: GFP-LC3B (green), Halo-B55α/δ or Halo-CDK16/17/18 (red), SNAP-WIPI2B (blue). Scale bar: 2 μm. (F) Representative micrographs of SNAP-WIPI2B in the distal axonal tip of DRG neurons from 24 mo mice ectopically expressing SNAP-WIPI2B and Halo-B55α (left) or Halo-CDK16 (right). Scale bar: 2 μm. (G) Quantification of SNAP-WIPI2B puncta in the distal axonal tip of GFP-LC3B DRG neurons ectopically expressing SNAP-WIPI2B and Halo-B55α, Halo-B55δ, Halo-CDK16, Halo-CDK17, or Halo-CDK18 (violin plot, median + first and third quartiles; n > 30 across three biological replicates for each condition). Unmarked comparisons are not significant (p > 0.05), ****p ≤ 0.0001, ***p ≤ 0.001 by Kruskal-Wallis test with Dunn’s multiple comparisons test. (H) Representative time-lapse micrographs of GFP-LC3B in the distal axonal tip of DRG neurons depicting an autophagosome biogenesis event (white arrowhead). Scale bar: 2 μm. (I) Quantification of the rate of AV biogenesis (GFP-LC3B puncta) in DRG neurons from 11 mo mice treated with control, B55α, B55δ, or CDK16 siRNA (violin plot, median + first and third quartiles; n > 30 across three biological replicates for each siRNA condition). **p ≤ 0.01 by Kruskal-Wallis test with Dunn’s multiple comparisons test.

We next asked whether ectopic expression of Halo-B55α/δ and Halo-CDK16/17/18 modulated formation of WIPI2B puncta. WIPI2B is recruited to the phagophore upon autophagy induction, forming punctate structures that colocalize with other early autophagy components (Stavoe et al., 2019; Polson et al., 2010). We previously found that expression of phospho-mimetic WIPI2B in DRG neurons prevented formation of autophagosomes and that phospho-mimetic WIPI2B localized to the phagophore half as long as phospho-dead WIPI2B (Stavoe et al., 2019). Thus, we hypothesized that overexpression of CDK16 would prevent formation of WIPI2B puncta in the DRG distal axon. Indeed, ectopic expression of Halo-CDK16 significantly diminished the number of SNAP-WIPI2B puncta in the DRG axonal tip as compared to Halo-B55α and Halo-B55δ ectopic expression (Fig. 6F, 6G). These data suggest that CDK16 regulates WIPI2B puncta formation.

We then asked how siRNA-mediated knockdown of B55α, B55δ, and CDK16 altered rates of autophagosome biogenesis. We first examined siRNA knockdown efficiency in murine HT-22 cells and verified knockdown of B55α, B55δ, and CDK16 by western blot. We found that all three siRNA pools were highly efficient: we achieved 77.5%, 61.9%, and 96.3% knockdown of B55α, B55δ, and CDK16, respectively (Fig. S2F-H). In primary neurons, as before, we used the GFP-LC3B reporter for autophagosomes to measure autophagosome biogenesis. We treated DRG neurons with control or targeted siRNA, and then used time-lapse spinning disk confocal microscopy to quantify the appearance of discrete GFP-LC3B puncta (autophagic vesicles, or AVs) at axonal distal tips within a ten-minute time window, as previously described (Fig. 6H) (Stavoe et al., 2019; Tsong et al., 2023). siRNA-mediated knockdown of B55α significantly decreased the rate of autophagosome biogenesis by approximately 50% (Fig. 6I). However, knockdown of B55δ did not alter the rate of autophagosome biogenesis (Fig. 6I), suggesting that B55α rather than B55δ is the dominant PP2A regulatory subunit involved in regulating autophagosome biogenesis in DRG neurons. These data also support our previous observation that loss of PP2A function impairs neuronal autophagy. Knockdown of CDK16 did not alter rates of autophagosome biogenesis (Fig. 6I), suggesting that there may be compensation or redundancy by CDK17 or CDK18.

Taken together, our data support a model in which PP2A-B55α dephosphorylates WIPI2B S395, promoting its function at the growing autophagosome. Then, CDK16 phosphorylates WIPI2B S395, to promote WIPI2B dissociation from the membrane, enabling expansion of the phagophore around cargo and maintenance of the cytosolic pool of WIPI2B for subsequent autophagosome formation. The proper recruitment of WIPI2B to and dissociation of WIPI2B from the phagophore is important for the proper formation of the autophagosome. Thus, PP2A and CDK16 work in opposition to regulate WIPI2B S395 phosphorylation, WIPI2B function, and autophagosome biogenesis in neurons.

## DISCUSSION

We previously identified WIPI2B S395 phosphorylation as a potential regulatory mechanism for WIPI2B function in autophagosome biogenesis (Stavoe et al., 2019). Here, we leveraged the genetic advantages of *C. elegans* as a screening platform to identify PP2A and CDK16 as regulators of WIPI2B S395 phosphorylation *in vivo.* We further validated these results by demonstrating direct enzymatic activity of mammalian recombinant PP2A and CDK16 on mammalian WIPI2B *in vitro*. In primary murine neurons, we observed colocalization of PP2A components and CDK16 with WIPI2B at LC3B-labelled autophagosomes. We also demonstrated that manipulating PP2A and CDK16 expression alters WIPI2B puncta formation and rates of autophagosome biogenesis in primary murine neurons. Thus, we have identified PP2A and CDK16 as regulators of neuronal autophagosome biogenesis.

Autophagy is a complex pathway that is tightly regulated in neurons. It is becoming increasingly clear that post-translational modifications are important regulatory mechanisms for autophagy. Such post-translational modifications modulate the subcellular localization of, interaction between, and activity of autophagy constituents (Xie et al., 2014). For example, phosphorylation of core autophagy proteins ULK1 and BECLIN1 modulates their function (Wong et al., 2015; Fujiwara et al., 2016). Identifying these post-translational modifications, understanding their effects, and elucidating the signaling cascades that govern them are essential for understanding the molecular mechanisms that regulate autophagy.

Here, we identified PP2A-B55α as a phosphatase that regulates WIPI2B S395 phosphorylation, whereby PP2A-B55α activity promotes neuronal autophagosome biogenesis. PP2A is a highly ubiquitous and promiscuous phosphatase, with specific substrates targeted for dephosphorylation by the PP2A regulatory subunits. PP2A-mediated dephosphorylation of autophagy proteins ULK1 and BECLIN1 has been previously described. In one study, the dephosphorylation of ULK1 by PP2A was found to enhance autophagic activity in mammalian immortalized cell lines (Wong et al., 2015), congruent with our findings here for PP2A regulation of WIPI2B S395. Conversely, a separate study found that dephosphorylation of BECLIN1 by PP2A inhibited autophagy (Fujiwara et al., 2016). Both studies identified B55α as the relevant PP2A regulatory subunit. It is interesting to note that the activity of the same phosphatase can have opposite effects on autophagy based on the targeted protein. Our data now adds to this intricate network of dephosphorylation, indicating that dephosphorylation of WIPI2B S395 by PP2A-B55α also promotes autophagosome biogenesis in neurons.

Our data identifies CDK16 as a kinase that opposes PP2A activity to regulate WIPI2B S395 phosphorylation. CDK16 (PCTAIRE1) along with CDK17 (PCTAIRE2) and CDK18 (PCTAIRE3) are members of an under-studied, highly conserved group of cyclin-dependent kinases. However, instead of regulating the cell cycle, CDK16 binds to Cyclin Y (Ou et al., 2010; Mikolcevic et al., 2012), a cyclin whose expression is not dependent on the cell cycle and is expressed in the mammalian brain (Cho et al., 2015; Joe et al., 2017). Furthermore, the PCTAIREs are highly expressed in post-mitotic tissues, including neurons. While the three members do share high sequence similarity, they have unique expression patterns and likely individual functions (Cole, 2009; de Oliveira Pepino et al., 2021). CDK16 has previously been implicated in autophagy regulation in immortalized cell lines, whereby phosphorylation of Cyclin Y by AMPK promotes CDK16 kinase activity and subsequent autophagy induction (Dohmen et al., 2020). The functions of CDK17 and CDK18 are less clear, but CDK18 has previously been implicated in tau phosphorylation (Herskovits and Davies, 2006). Here we identify a role for CDK16 in regulating WIPI2B function in neurons. Similar to the intricate regulatory effects of PP2A-B55α dephosphorylation on autophagy, our data now add to the complexity of CDK16 modulation of autophagy.

Previous reports identified mTORC1 as a kinase that regulates WIPI2B S395 phosphorylation (Wan et al., 2018; Hsu et al., 2011). However, these studies were conducted in immortalized cell lines, not *in vivo* or in neural tissues. mTORC1 is a master regulator of autophagy in immortalized cell lines, with starvation or inhibition of mTORC1 causing massive upregulation of autophagy (Abada and Elazar, 2014; Hale et al., 2013). In contrast, neuronal autophagy is not responsive to mTORC1 inhibition or other “traditional” activators of autophagy in primary mammalian neurons (Tsvetkov et al., 2010; Maday and Holzbaur, 2016). We did note that inhibition of mTORC1 by Torin 1 did modestly decrease WIPI2B S395 phosphorylation by whole brain homogenate (Fig. 5A, 5B), suggesting that mTORC1 may have a limited capability to phosphorylate WIPI2B S395.

Upregulating neuronal autophagy during aging or in the context of neurodegenerative disease has been a long-standing therapeutic goal, since increasing autophagy function in these contexts is hypothesized to support the turnover of damaged organelles and prevent or slow the build-up of protein aggregates to preserve neuronal health. Many studies have attempted to increase neuronal autophagy by inhibiting mTORC1, which triggers autophagy induction in immortalized cell lines (Abada and Elazar, 2014; Hale et al., 2013). However, both *in vitro* and *in vivo* studies were largely unsuccessful (Tsvetkov et al., 2010; Maday and Holzbaur, 2016; Mizushima et al., 2004; Fox et al., 2010). Additionally, our previous work demonstrates that the autophagy induction complex is unaffected by aging. In DRG neurons from aged mice, the induction and nucleation complex proteins were successfully recruited to the phagophore. Instead, we showed that ectopic expression of WIPI2B restores autophagosome biogenesis in aged neurons (Stavoe et al., 2019). As such, WIPI2B is an attractive target to increase rates of neuronal autophagosome biogenesis during aging. Since WIPI2B has no apparent enzymatic activity itself, it is refractory to therapeutic targeting. However, as WIPI2B S395 phosphorylation regulates its function, the molecular pathways that regulate WIPI2B S395 phosphorylation present more promising therapeutic targets to increase autophagosome formation, thereby supporting neuronal health during aging. Thus, we have now identified two promising therapeutic targets to modulate neuronal autophagy: PP2A-B55α and CDK16.

Importantly, while a certain degree of autophagy upregulation may prove beneficial, overactive autophagy is also implicated in some disease states (Sridhar et al., 2012). Overactive autophagy can lead to aberrant degradation and strain the cell due to excessive use of resources for the autophagy process. Thus, autophagy is a double-edged sword and maintaining an optimal level of autophagy should be the therapeutic goal. We note that the PVD axon outgrowth phenotype was not as drastic in phospho-mimetic *atg-18(S375E)* mutant worms as in *atg-18(LOF)* or *atg-18(FTTG)* mutant animals (Fig. 1C). This “in-between” phenotype may reflect a fine-tuning aspect of WIPI2B phosphorylation in regulating neuronal autophagy. Targeting a fine-tuning regulatory mechanism such as WIPI2B phosphorylation may provide therapeutically beneficial upregulation without surpassing certain thresholds that can lead to detrimental effects. Thus, WIPI2B phosphorylation regulators may present a uniquely optimal therapeutic target for achieving “Goldilocks” levels of autophagosome biogenesis in aged neurons.

## MATERIALS AND METHODS

### Reagents

#### C. elegans strains and maintenance

*C. elegans* strains were maintained at room temperature on NGM plates seeded with OP50 *E. Coli*. The N2 Bristol strain was used as the wild-type control and was obtained from the *Caenorhabditis* Genetics Center (CGC). Other strains obtained from the CGC include: VC30247 *atg-18(gk447096),* RB1667 *tax-6(ok2065),* EU1062 *sur-6(or550),* JH2787 *pptr-1(tm3103),* RB1471 *pct-1(ok1707),* VC8 *jnk-1(gk7),* KU25 *pmk-1(km25),* FJ1519 *cdk-5(gm336),* WM99 *cdk-1(ne2257),* and CB408 *unc-43(e408)*. Information about the specific alleles can be found in Supplementary Table S2. Strains obtained from the CGC were genotyped via PCR and restriction digest (if applicable) upon receipt. The CX9797 *kyIs445* [des-2p::mCh::rab-3; des-2p::sad-1::gfp; odr-1p::DsRed] strain was obtained from the Daniel Colón-Ramos at Yale University.

#### CRISPR-generated transgenic lines

An adapted co-CRISPR protocol (Ghanta and Mello, 2020) was used to generate VOE192 *kyIs445; atg-18(zny7[S375E]),* VOE201 *kyIs445; atg-18(zny6[S375A]),* VOE246 *kyIs445; atg-18(zny10[FTTG]),* and VOE489 *kyIs445; pptr-2(zny27[Y220*])*. For the co-CRISPR marker, the *dpy-10(cn64)* lesion, which produces worms with the roller or dumpy phenotype, was recapitulated (Dickinson and Goldstein, 2016). All CRISPR reagents were designed with and obtained from Integrated DNA Technologies (IDT). CRISPR was performed in CX9797 *kyIs445* worms due to the proximity of the genetic loci of *kyIs445*, *atg-18,* and *pptr-2*. CRISPR edits were confirmed through Sanger sequencing. Successful lines were then outcrossed five times to N2 animals and confirmed to have no off-target edits by the CRISPR guide RNA and co-CRISPR guide RNA (as predicted by IDT) through Sanger sequencing before use in the study.

#### Mice

GFP-LC3B transgenic mice (strain: B6.Cg-Tg(CAG-EGFP/LC3)53Nmi/NmiRbrc) originally generated by N. Mizushima (Tokyo Medical and Dental University, Tokyo, Japan) were obtained from RIKEN BioResource Center in Japan (RBRC00806). These mice were bred with C57BL/6J mice obtained from The Jackson Laboratory (000664). Hemizygous mice were used in experiments.

#### Plasmids

Plasmid constructs used in this study include Halo-B55α (subcloned from Addgene 13804), Halo-B55δ (cloned by PCR from whole mouse brain cDNA), Halo-CDK16 (subcloned from Addgene 1999), Halo-CDK17 (subcloned from Addgene 23845), Halo-CDK18 (subcloned from Addgene 23710), and SNAP-WIPI2B (Stavoe et al., 2019). All Halo-tagged constructs were cloned into pHTN (Promega). We note that when CDK16 was cloned into pHTC (Promega), the CDK16-Halo construct was refractory to expression in primary DRG neurons. All plasmids were confirmed with Sanger or whole-plasmid sequencing.

### Primary neuron culture

Mice were euthanized prior to dissection. All animal protocols were approved by the Institutional Animal Care and Use Committee at UTHealth Houston. DRG neurons were isolated as previously described (Perlson et al., 2009) from 11 mo or 24 mo mice of either sex. DRG neurons were plated on glass-bottomed dishes (MatTek Corporation, P35G-1.5-14-C) and cultured in F-12 Ham’s media (Gibco, 11765-047) with 10% heat-inactivated FBS (HyCLone, SH30071.03), 100 U/mL penicillin, and 100 μg/mL streptomycin (Gibco, 15140122). DRG neurons were imaged after being maintained for 2 days at 37°C in a 5% CO_2_ incubator.

Neurons were transfected with plasmid DNA or siRNA using a 4D-Nucleofector (Lonza) as per the manufacturer’s instructions before plating. For ectopic expression experiments, neurons were nucleofected with 0.25 μg of Halo-tagged constructs and 0.25 μg of SNAP-WIPI2B. Prior to imaging, DRG neurons were incubated with 100 nM Halo-TMR (HHMI, Janelia) and 100 nM SNAP-Cell 647 SiR (New England Biolabs) ligand for at least 30 min at 37°C in a 5% CO_2_ incubator. After incubation, neurons were washed three times with complete equilibrated F-12 media, with the final wash remaining on the neurons for at least 15 min at 37°C in a 5% CO_2_ incubator. For knockdown experiments, siRNAs were diluted to 50 pmol/μL with RNAse-free water and 5X siRNA Buffer (Dharmacon/Horizon Discovery, B-0020000-UB-100). Neurons were nucleofected with 100 pmol of non-targeting (Dharmacon/Horizon Discover, D-001810-0105), B55α (Dharmacon/Horizon Discovery, L-047957-00-0005), B55δ (Dharmacon/Horizon Discovery, L-057039-02-0005), or CDK16 (Dharmacon/Horizon Discovery, L-040144-00-0010) siRNA in INB buffer with 5 pmol of Cy5-labelled non-targeting siRNA (Dharmacon/Horizon Discovery).

Prior to use in DRG neurons, siRNA knockdown was validated in murine HT-22 cells that were cultured in 6-well plates in DMEM with 10% FBS and Pen/Strep and maintained at 37°C in a 5% CO_2_ incubator. Briefly, HT-22 cells were transfected with experimental or control siRNA (see above) with RNAiMax (Invitrogen), following the manufacturer’s instructions. 150 pmol of each siRNA was added per well. Transfected cells were incubated for 2 days. HT-22 cells were then harvested in RIPA buffer; clarified lysates were denatured and boiled in 4X denaturing buffer (recipe from Li-Cor).

### Live imaging and image analysis

#### PVD axon imaging and quantification

Worms were mounted onto 2% agarose pads and immobilized in 10 mM levamisole (MilliporeSigma) prior to imaging. Z-stacks (step size: 0.9 μm) were acquired on an upright epifluorescent microscope (Nikon Instruments) with an CFI60 Plan Apochromat Lambda 20X objective and captured with a sCMOS camera (Excelitas) using the Nikon Elements software (Nikon Instruments).

All measurements were performed on maximum intensity projections of the acquired z-stacks using FIJI. PVD axons were measured using the mCh::rab-3 marker for presynapses along the ventral nerve cord, not including the distance between the ventral nerve cord and the cell body. Body length was measured from the tip of the fluorescent signal in the head to the fluorescent signal in the tail. Measured PVD lengths were normalized to the average body length for each genotype and then normalized to the wild-type mean in Microsoft Excel.

#### Live-cell imaging of DRGs and quantification

DRG neurons were switched to Hibernate A low fluorescence nutrient media (BrainBits, HALF500) with 2% B27 (Gibco, A3582801) and 2mM GlutaMAX (Gibco, 35050061) for microscopy. Confocal images were captured at 37°C using a spinning-disk confocal (Nikon Ti2 Inverted Confocal with Yokogawa W1 Spinning Disk Package) with an Apochromat 100x, 1.4 NA oil immersion objective (Nikon Instruments) and fitted with an environmental chamber. Digital micrographs were acquired with a back-illuminated cCMOS camera (Teledyne Photometrics) using Nikon Elements software.

Time-lapse images were acquired for 10 min with a frame every 3s. The Perfect Focus System was used to maintain Z position during time-lapse acquisition. Multiple channels were acquired consecutively, with the green (488 nm) channel captured first, followed by red (561 nm), and far-red (640 nm). DRG neurons were imaged at the distal axon tips and were selected for imaging based on morphological criteria and low expression of transfected constructs.

All image analysis was performed on raw data with FIJI with the brightness and contrast adjusted equally across all images within a series. Autophagosome biogenesis was quantified as previously described (Stavoe et al., 2019; Tsong et al., 2023). Briefly, GFP-LC3B puncta were tracked manually, with a biogenesis event defined as the *de novo* appearance of a GFP-LC3B puncta that increased in fluorescence intensity within the imaging window. GFP-LC3B puncta already present at the start of imaging were not counted. SNAP-WIPI2B puncta were quantified by manually counting the already present and newly generated puncta within the time window. Colocalization was quantified using plot profiles and normalized to the max fluorescence signal for each captured channel.

### in vitro assays

#### Generation of WIPI2B tail substrate

For the in vitro assays, a GST- HA-tagged peptide containing the last 78 amino acids of human WIPI2B (WIPI2B tail) was produced by cloning the fragment into the pGEX4T-1 bacterial expression plasmid. Protein expression was accomplished by transforming this plasmid along with a second bacterial plasmid expressing lambda phosphatase (Addgene 79748) into BL21(DE3) *E. Coli* competent cells (Invitrogen). Selection of dual transformed colonies was performed on LB plates containing 100 μg/mL spectinomycin (MP Biomedicals) and 50 μg/mL ampicillin (Thermo Fisher). Protein expression was accomplished by splitting overnight cultures 1:50 in fresh LB with 100 μg/mL spectinomycin and 50 μg/mL ampicillin, grown to OD_600_=0.6, and then protein was induced by adding 1mM IPTG and shaking for 4 hrs at 37°C. Bacteria were harvested by centrifugation at 5000 x *g* for 15 min at 4°C and pellets were frozen at -20°C until use. The bacterial pellet was lysed in B-PER Bacterial Protein Extraction Reagent (Thermo Fisher) supplemented with 1X cOmplete^TM^ Mini protease inhibitor cocktail (Roche), 0.1 μg/μL lysozyme (Thermo Fisher), and 5 U/mL DNase I (Thermo Fisher). The clarified supernatant was then aliquoted and frozen at -20°C. Prior to use in the assays, the aliquoted WIPI2B tail was purified with Glutathione Sepharose 4B (Cytiva) affinity chromatography resin. As a final step, the purified WIPI2B tail was washed three times with 1X TBS to remove all protease inhibitors and resuspended in 1X TBS before use in the assays as described below.

#### Expression and purification of CaMKIIα and calmodulin

Expression and purification of CaMKIIα was accomplished using a baculovirus system as previously described (Putkey and Waxham, 1996; Gaertner et al., 2004). Following purification using CaM-Sepharose affinity chromatography, purified protein was dialyzed into 20 mM HEPES, 0.5M NaCl, and 10% glycerol, pH7.4 and stored as single use aliquots frozen at -80°C. CaMKIIα protein was quantified using a calculated extinction coefficient at 280 nm of 1.23 = 1 mg/mL. Calmodulin was expressed from a codon optimized cDNA in the pET-23d plasmid expressed in BL21(DE3) *E. coli* and purified as previously described (Putkey and Waxham, 1996). Briefly, the pellets were resuspended in 50 mM Tris, pH 7.5, and 1 mM EDTA, and sonicated before centrifugation at 18,000 x *g* for 1 hr at 4°C. The supernatant was collected and applied to a phenyl-Sepharose CL-4B column (GE Healthcare) previously equilibrated with 50 mM Tris, pH 7.5 and 2 mM CaCl_2._ After sample application, the column was first washed with 50 mM Tris, pH 7.5, 2 mM CaCl_2_ + 0.5 M NaCl and then with 50 mM Tris, pH 7.5 and 2 mM CaCl_2._ Calmodulin was eluted with 50 mM Tris, pH 7.5, and 10 mM EGTA. The protein was extensively dialyzed into 50 mM MOPS, pH 7.0 and stored in this buffer as aliquots at -80°C. The concentration of each protein was quantified by using the extinction coefficient of A_276_ = 2560 M^-1^cm^-1^.

#### Dephosphorylation assay

A mouse brain was isolated and homogenized in RIPA buffer [1X PBS, 10 mM Na_2_HPO_4_, 2 mM KH_2_PO_4_ pH 7.4, 1% Triton X-100, 0.5% deoxycholate, 0.1% SDS, and 1X cOmplete^TM^ Mini protease inhibitor cocktail]. The homogenate was then aliquoted, snap frozen in liquid nitrogen, and stored at -80°C. Phosphorylation of the purified WIPI2B tail was achieved by incubating the purified WIPI2B tail with ∼10 μg brain homogenate and 200 μM ATP (Cell Signaling Technology) in 1X kinase buffer [25 mM Tris-HCl (pH 7.5), 5 mM beta-glycerophosphate, 2 mM dithiothreitol (DTT), 0.1 mM Na_3_VO_4_, 10 mM MgCl_2_] (Cell Signaling Technology) in a total volume of 50 μL at 30°C for 30 min. Following this treatment, the WIPI2B tail was washed two times with 0.05% TBS + Tween-20 and then two times with 1X TBS to remove the brain homogenate. The phosphorylated purified WIPI2B tail was then incubated with 0.4 μg of active recombinant human PP2A (Abcam) in phosphatase buffer [40 mM Tris-HCl, 34 mM MgCl_2_, 4 mM EDTA, 2 mM DTT, 0.05 mg/mL BSA] in a total volume of 50 μL at 37°C for 1 hr.

#### Phosphorylation assay

For the *in vitro* phosphorylation assay with kinase inhibitors, purified WIPI2B tail was incubated with ∼10 μg brain homogenate, 200 μM ATP, and the indicated kinase inhibitor (400 nM Cdk1/2 inhibitor III (MilliporeSigma), 1 μM Torin 1 (Cell Signaling Technology), 1 μM autocamtide-2-related inhibitory peptide (AIP) (Tocris)) in 1X kinase buffer in a total volume of 50 μL at 30°C for 30 min. For the *in vitro* phosphorylation assay with CDK16, purified WIPI2B tail was incubated with 0.45 μg of active recombinant human CDK16 + Cyclin Y (Abcam) in 1X kinase buffer with or without 200 μM ATP and 400 nM Cdk1/2 inhibitor III in a total volume of 50 μL at 30°C for 30 min. For the *in vitro* phosphorylation assay with CaMKIIα, the purified WIPI2B tail was incubated with 100 ng of recombinant CaMKIIα along with 1 μM calmodulin, 100 μM Ca^2+^, and 200 μM ATP in 1X kinase buffer in a total volume of 50 μL at 30°C for 30 min.

All reactions were stopped by adding 4X denaturing buffer. Samples were then boiled at 95°C for 5 min and stored at -20°C. All assays were performed in triplicate.

### Immunoblotting

*In vitro* assay samples and HT-22 cell lysates were analyzed via SDS-PAGE immunoblot transferred onto FL PVDF membranes (MilliporeSigma, IPFL00010) and visualized with fluorescent secondary antibodies using an Odyssey® CLx imaging system (Li-Cor). See Key Resources Table for antibodies used. All western blots were analyzed with Image Studio (Li-Cor). For the HT-22 cell lysates, total protein was used as a loading control (REVERT™ Total Protein Stain, Li-Cor). The normalization factor is listed below the blot as a percent.

### Statistics

Data were assembled in Microsoft Excel and Prism 10 (GraphPad) was used to generate graphs and perform statistical tests. Data were tested for normality by the D’Agostino and Pearson test. The appropriate statistical test was then used based on this analysis of normality. Specific statistical tests are indicated in the corresponding figure legends.

## KEY RESOURCES TABLE

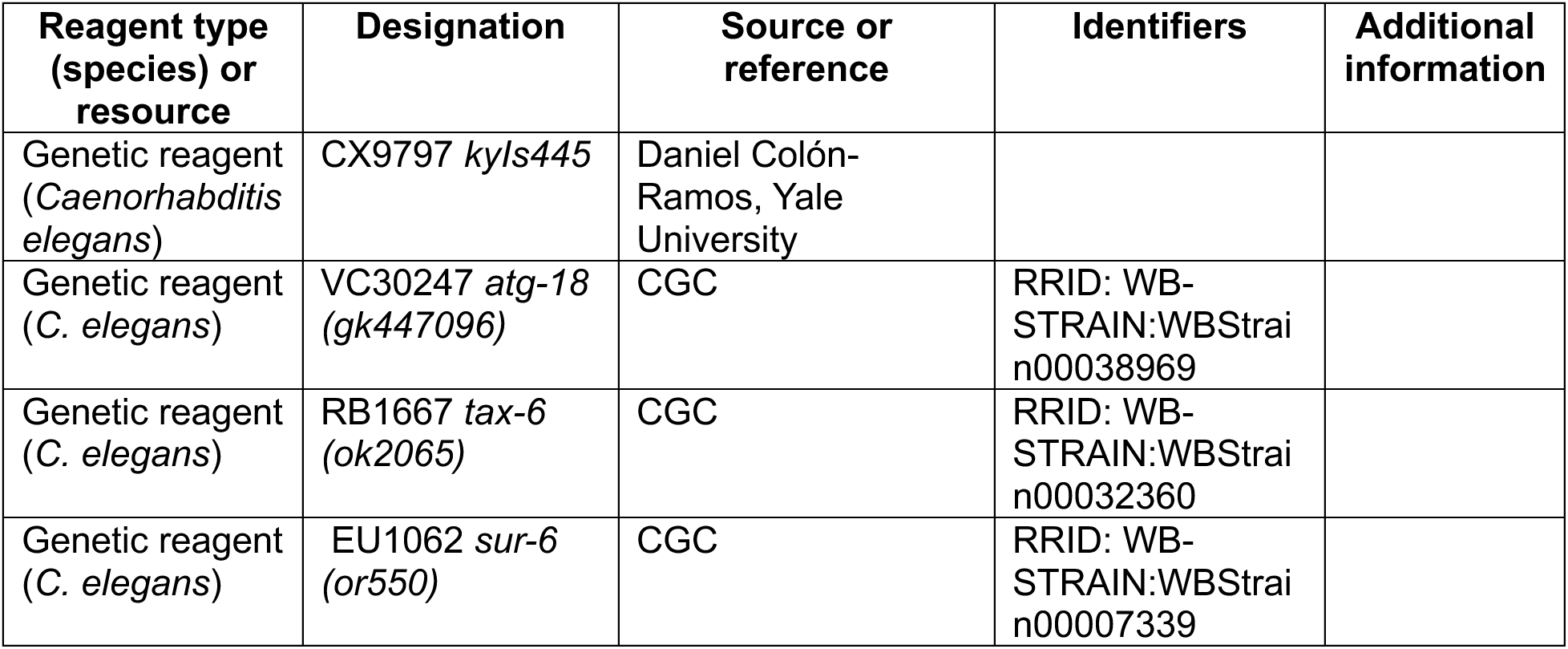
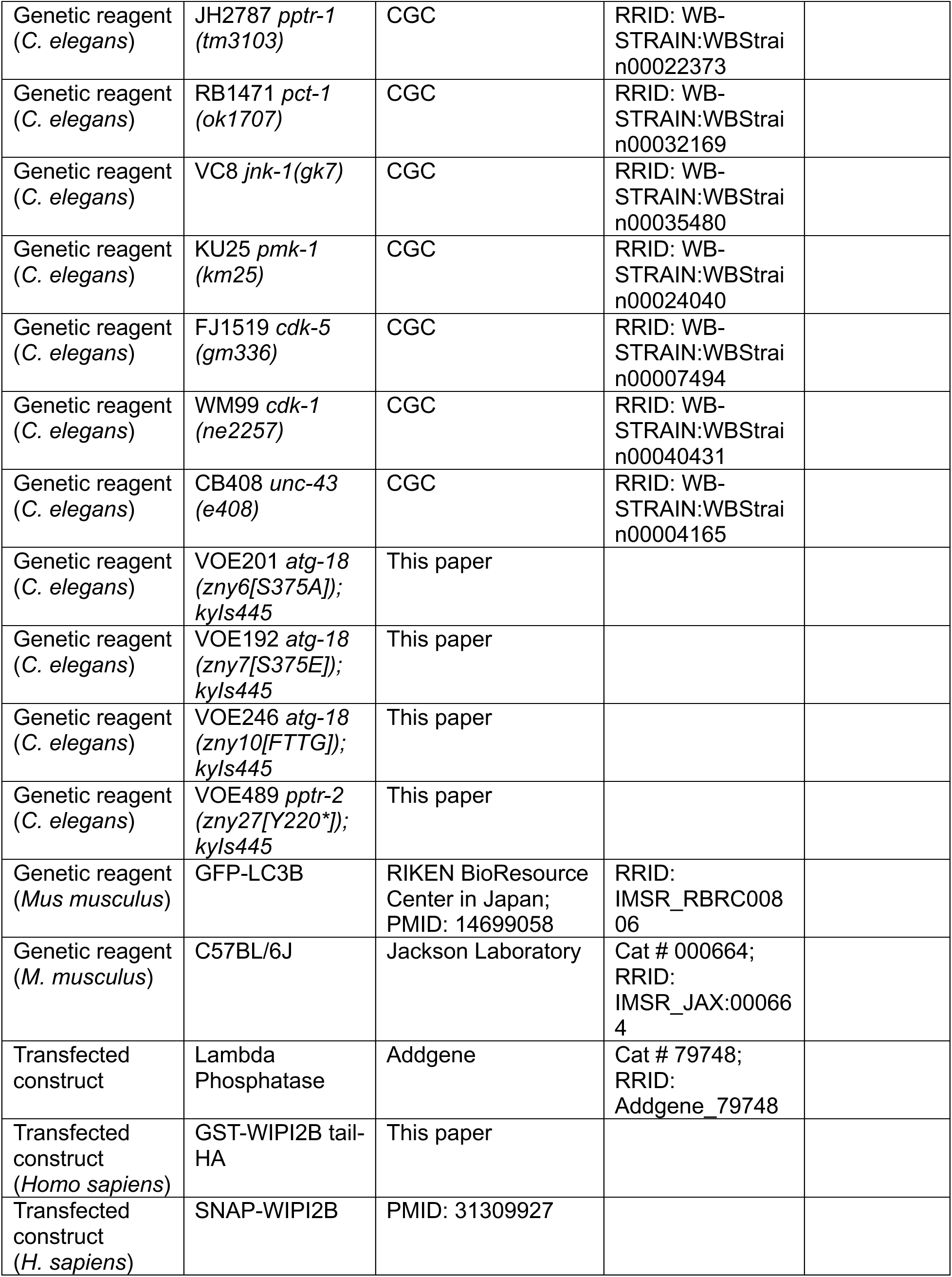
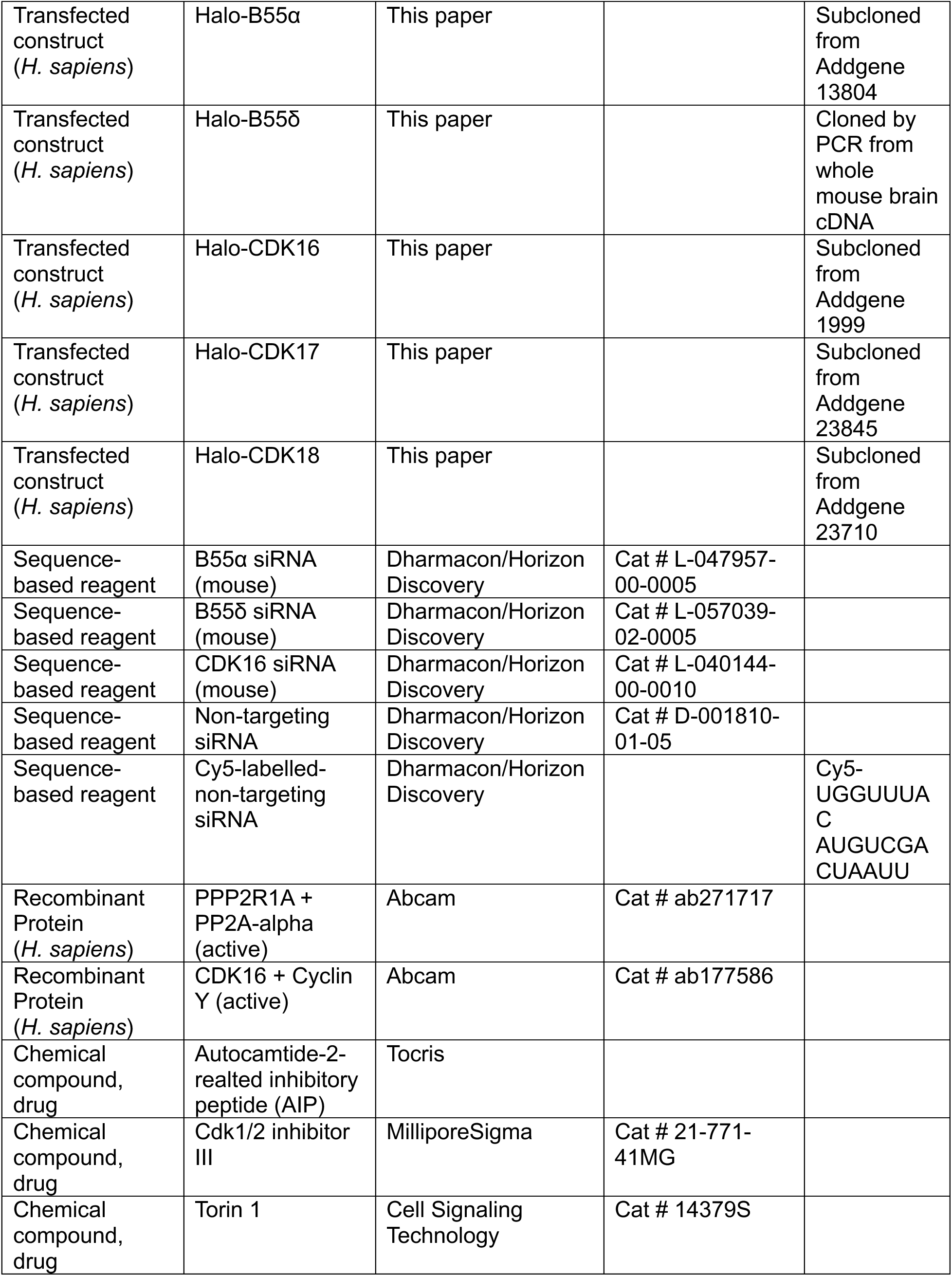
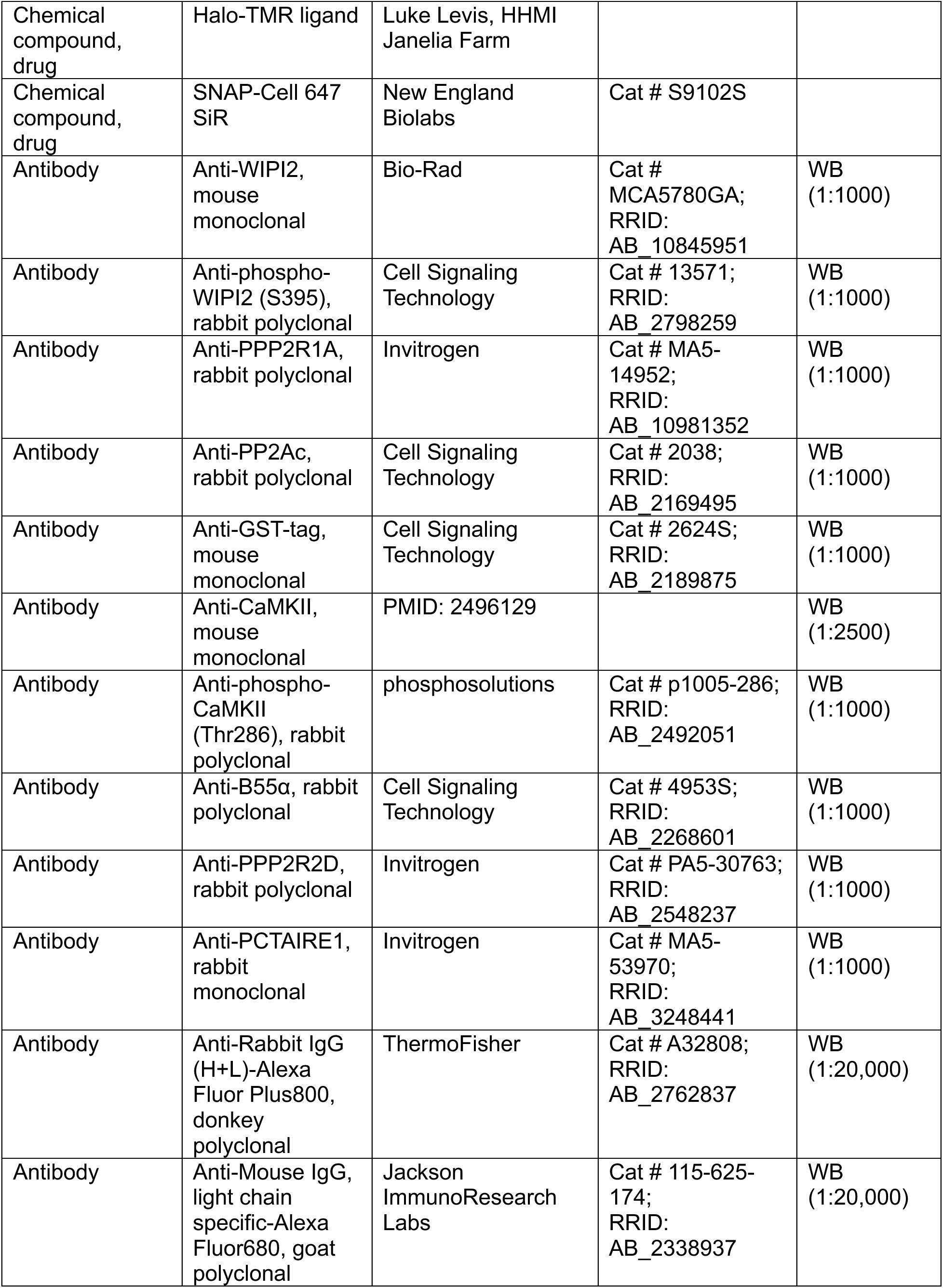
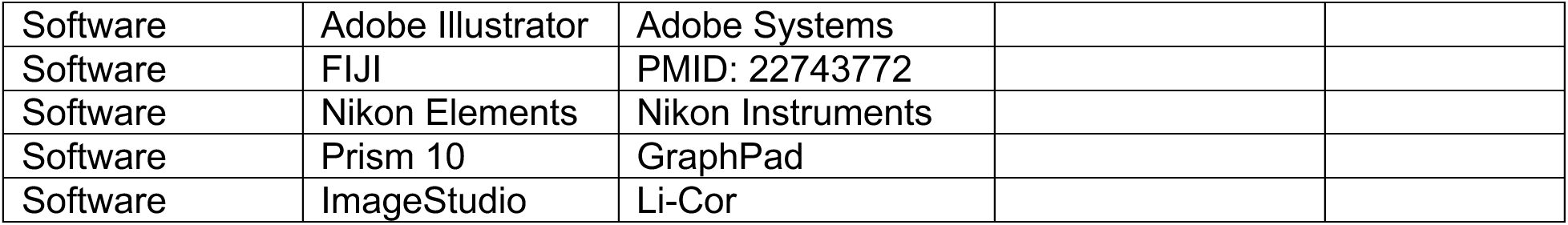

## SUPPLEMENTAL MATERIAL

Supplementary Table 1 lists fifty proteins identified as potential kinases for WIPI2B S395 by Johnson et al., 2023 and by a kinase prediction algorithm (PhosphoNET). Supplementary Table 2 lists the *C. elegans* alleles used in this study. The table includes the *C. elegans* gene name, the mammalian ortholog(s), and the specific allele and lesion information. Fig. S1 presents data showing that *atg-18* is not epistatic to *cdk-5* or *unc-43* in regulating PVD axon length. Fig. S2 shows examples of colocalization events in which Halo-B55α, Halo-B55δ, Halo-CDK16, Halo-CDK17, or Halo-CDK18 colocalized with SNAP-WIPI2B but not with GFP-LC3B. Fig. S2 also shows the validation of the siRNA-mediated knockdown of B55α, B55δ, and CDK16 in HT-22 cells.

## ACKNOWLEDGEMENTS

The authors acknowledge the technical assistance of past and current Stavoe lab members Sylvia LeBlanc, Beatriz Rios, Andrew Martinez, and Dheerj Jasuja. We would also like to acknowledge Dr. Daniel Colon-Ramos for providing a key *C. elegans* strain and Dr. Rachel Arey and her lab for helpful discussions.

## ABBREVIATIONS

DRG: dorsal root ganglia
LOF: loss-of-function
PI3P: phosphatidylinositol 3-phosphate
PP2A: protein phosphatase 2A

## SUPPLEMENTARY FIGURE LEGENDS

Table S1. List of candidate kinases

List of top 50 kinases that were predicted to regulate WIPI2B S395 phosphorylation as identified by Johnson et al., 2023 (column 3) and by the prediction algorithm, PhosphoNET (column 4). Kinases are ordered based on the ranking determined by Johnson et al., 2023. Asterisks denote kinases that were examined in this study.

Table S2. List of *C. elegans* genes and mammalian orthologs with allele information

List of all *C. elegans* genes analyzed in this paper with the corresponding mammalian ortholog (column 2), examined alleles (column 3), nature of genetic lesion (column 4), and effect on the protein (column 5). Rows in orange are genes encoding phosphatase and rows in blue are genes encoding kinases.

Figure S1. *atg-18* is not epistatic to *cdk-5* or *unc-43* (A) Quantification of normalized PVD axon length for epistatic analysis of *unc-43* and *atg-18* in mutant L4 animals (violin plot, median + first and third quartiles; n > 100 animals across three replicates). ****p ≤ 0.0001, **p ≤ 0.01, *p ≤ 0.05, ns p > 0.05 by Kruskal-Wallis test with Dunn’s multiple comparisons test. (B) Quantification of normalized PVD axon length for epistatic analysis of *cdk-5* and *atg-18* in mutant L4 animals (violin plot, median + first and third quartiles; n > 100 animals across three replicates). ****p ≤ 0.0001, **p ≤ 0.01, *p ≤ 0.05, ns p > 0.05 by Kruskal-Wallis test with Dunn’s multiple comparisons test. (C-H) Representative images of PVD axon presynapses (visualized with mCh::rab-3) in wild-type (C), *atg-18(S375E)* (D), *unc-43* (E), *unc-43; atg-18(S375E)* (F), *cdk-5* (G), *cdk-5; atg-18(S375E)* (H) L4 animals. Dashed bracket indicates the measured axon. Scale bar: 20 μm.

Figure S2. Ectopically expressed PP2A components and CDK16/17/18 co-localize with WIPI2B in DRG neurons and validation of siRNA mediated knockdown in HT-22 cells (A-E) Representative merged micrographs of distal axonal tips of GFP-LC3B DRG neurons from 22 mo mice ectopically expressing SNAP-WIPI2B and Halo-B55α (A), Halo-Β55δ (B), Halo-CDK16 (C), Halo-CDK17 (D), or Halo-CDK18 (E). Purple arrowheads indicate a colocalization event between Halo-constructs and SNAP-WIPI2B that are depicted in the magnified views and quantified in the corresponding line scan plot below the full micrograph. Borders of the magnified views and color of lines in the line scan plot denote the channel: GFP-LC3B (green), Halo-B55α/δ or Halo-CDK16/17/18 (red), SNAP-WIPI2B (blue). Scale bar: 2 μm. (F) Representative immunoblot and corresponding quantification (mean ± SEM, n = 3) of siRNA-mediated knockdown of B55α in HT-22 cells. Total protein was used as a loading control and for normalization during quantification (normalization factor indicated below each blot as a percentage). ***p ≤ 0.001 by one-way ANOVA with Tukey’s multiple comparisons test. (G) Representative immunoblot and corresponding quantification (mean ± SEM, n = 3) of siRNA-mediated knockdown of B55δ in HT-22 cells. Total protein was used as a loading control and for normalization during quantification (normalization factor indicated below each blot as a percentage). **p ≤ 0.01 by one-way ANOVA with Tukey’s multiple comparisons test. (H) Representative immunoblot and corresponding quantification (mean ± SEM, n = 3) of siRNA-mediated knockdown of CDK-16 in HT-22 cells. Total protein was used as a loading control and for normalization during quantification (normalization factor indicated below each blot as a percentage). ****p ≤ 0.0001 by one-way ANOVA with Tukey’s multiple comparisons test.

