## Supplemental Files for "PP2A and CDK16 antagonistically regulate WIPI2B phosphorylation and neuronal autophagosome biogenesis"

Table S1

| | Human Kinase Name | Human UniProt ID | Johnson et al., 2023 Rank | PhosphoNET Rank |
|---|---|---|---|---|
| | CDK6 | Q00534 | 1 | 28 |
| | CDK13 | Q14004 | 2 | 18 |
| | CDK4 | P11802 | 3 | 33 |
| | CDK12 | Q9NYV4 | 4 | 21 |
| | CDK9 | P50750 | 5 | 37 |
| | CDK2 | P24941 | 6 | 6 |
| | CDK8 | P49336 | 7 | |
| * | CDK1 | P06493 | 8 | 1 |
| | ERK1 | P27361 | 9 | 7 |
| * | CDK5 | Q00535 | 10 | 14 |
| | CDK19 | Q9BWU1 | 11 | 50 |
| | CDK10 | Q15131 | 12 | 23 |
| | ERK2 | P28482 | 13 | 8 |
| * | P38A (MAPK14) | Q16539 | 14 | 10 |
| | CDK7 | P50613 | 15 | 27 |
| | CDKL5 | O76039 | 16 | 9 |
| | NLK | Q9UBE8 | 17 | 38 |
| | MPSK1 | O75716 | 18 | |
| * | P38B (MAPK11) | Q15759 | 19 | 11 |
| | CDK3 | Q00526 | 20 | 4 |
| | ERK7 | Q8TD08 | 21 | 16 |
| | MOK | Q9UQ07 | 22 | 34 |
| | P38G | P53778 | 23 | 15 |
| | ICK | Q9UPZ9 | 24 | 29 |
| | P38D | O15264 | 25 | 25 |
| | SRPK3 | Q9UPE1 | 26 | |
| | MAK | P20794 | 27 | 30 |
| | ERK5 | Q13164 | 28 | 13 |
| * | CDK17 (PCTAIRE2) | Q00537 | 29 | 17 |
| | DYRK1A | Q13627 | 30 | |
| | CDKL1 | Q00532 | 31 | 20 |
| | HIPK3 | Q9H422 | 32 | |
| | CDK14 | O94921 | 33 | 31 |
| * | JNK2 (MAPK9) | P45984 | 34 | 5 |
| * | JNK3 (MAPK10) | P53779 | 35 | 3 |
| | SRPK1 | Q96SB4 | 36 | |
| | MTOR | P42345 | 37 | 35 |
| | HIPK4 | Q8NE63 | 38 | |
| * | CDK16 (PCTAIRE1) | Q04735 | 39 | 24 |

| Human Kinase Name | Human UniProt ID | Johnson et al., 2023 Rank | PhosphoNET Rank |
|---|---|---|---|
| SRPK2 | P78362 | 40 | |
| DYRK3 | O43781 | 41 | 47 |
| DYRK2 | Q92630 | 42 | 44 |
| * JNK1 (MAKP8) | P45983 | 43 | 2 |
| * CDK18 (PCTAIRE3) | Q07002 | 44 | 19 |
| DYRK1B | Q9Y463 | 45 | |
| HASPIN | Q8TF76 | 46 | |
| CLK3 | P49761 | 47 | |
| HIPK2 | Q9H2X6 | 48 | |
| PRP4 | Q13523 | 49 | 43 |
| KIS | Q8TAS1 | 50 | |

### Table S2

| *C. elegans* Gene | Mammalian Gene | Alleles | Lesion | Protein Effect |
|---|---|---|---|---|
| *atg-18* | Wipi1, Wipi2 | gk447069 | Q79 to amber stop | putative null |
| | | zny6 | S375A | S375 phosphorylation dead |
| | | zny7 | S375E | S375 phosphorylation mimic |
| | | zny10 | R228T, R229T | PI3P-binding deficient |
| *tax-6* | Ppp3cb, Ppp3cc | ok2065 | insertion/deletion | putative null |
| *sur-6* | Ppp2r2a, Ppp2r2d | or550 | W140R | hypomorph |
| *pptr-1* | Ppp2r5e | tm3103 | deletion | putative null |
| *pptr-2* | Ppp2r5d | zny27 | Y220 to stop | putative null |
| *pct-1* | Cdk16 | ok1707 | deletion | putative null |
| *jnk-1* | Mapk8, Mapk9, Mapk10 | gk7 | deletion | putative null |
| *pmk-1* | Mapk11, Mapk14 | km25 | deletion | putative null |
| *cdk-5* | Cdk5 | gm336 | deletion | putative null |
| *cdk-1* | Cdk1 | ne2257 | I173F | hypomorph |
| *unc-43* | Camk2d | e408 | S179L | hypomorph |

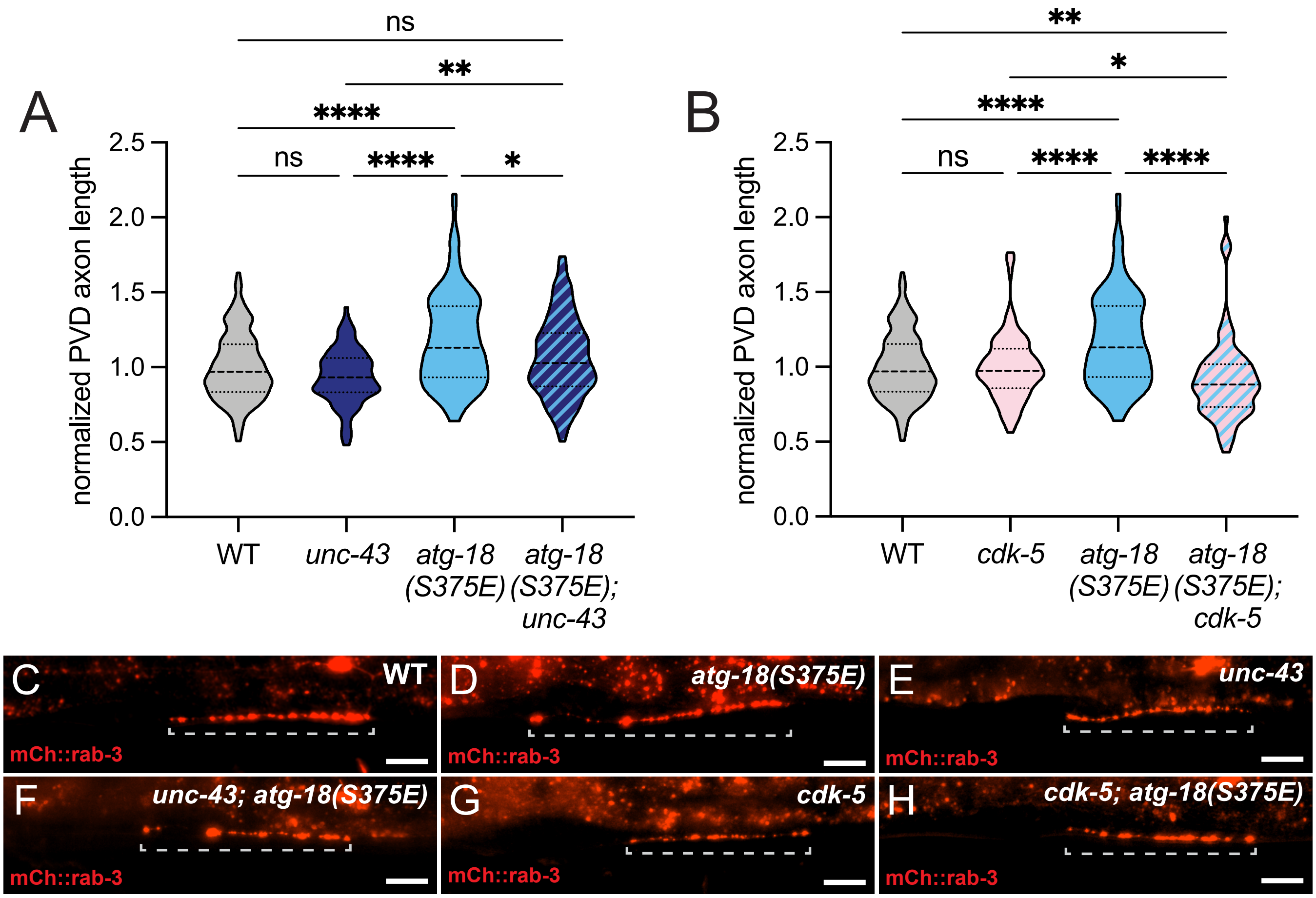
A
B
normalized PVD axon length
2.5
2.0
1.5
1.0
0.5
0.0
ns
**
****
ns
****
*
WT
unc-43
atg-18 (S375E)
atg-18 (S375E); unc-43
**
*
****
ns
****
****
cdk-5
atg-18 (S375E); cdk-5
C
WT
mCh::rab-3
D
atg-18(S375E)
E
unc-43
F
unc-43; atg-18(S375E)
G
cdk-5
H
cdk-5; atg-18(S375E)

Figure S1

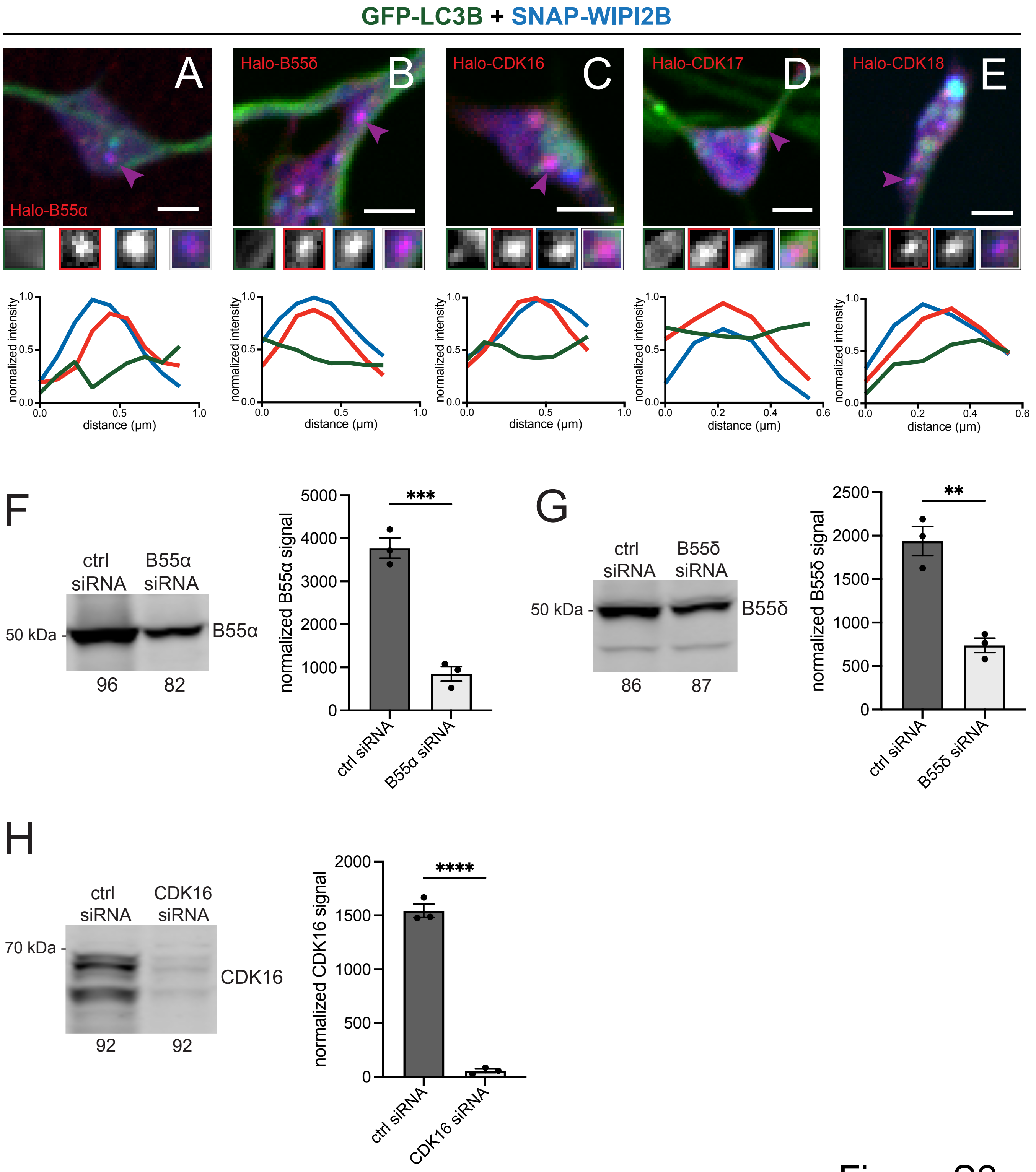

GFP-LC3B + SNAP-WIPI2B
A
Halo-B55α
B
Halo-B55δ
C
Halo-CDK16
D
Halo-CDK17
E
Halo-CDK18
normalized intensity
distance (µm)
F
ctrl siRNA
B55α siRNA
50 kDa
B55α
96
82
normalized B55α signal
***
ctrl siRNA
B55α siRNA
G
ctrl siRNA
B55δ siRNA
50 kDa
B55δ
86
87
normalized B55δ signal
**
ctrl siRNA
B55δ siRNA
H
ctrl siRNA
CDK16 siRNA
70 kDa
CDK16
92
92
normalized CDK16 signal
****
ctrl siRNA
CDK16 siRNA


Figure S2
